# Time- and lineage-resolved transcriptional profiling uncovers gene expression programs and clonal relationships that underlie human T lineage specification

**DOI:** 10.1101/2023.10.06.561277

**Authors:** Yale S. Michaels, Matthew C. Major, Becca Bonham-Carter, Jingqi Zhang, Tiam Heydari, John M. Edgar, Laura Greenstreet, Roser Vilarrasa-Blasi, Seungjoon Kim, Elizabeth L. Castle, Aden Forrow, M Iliana Ibanez-Rios, Carla Zimmerman, Yvonne Chung, Tara Stach, Nico Werschler, David JHF Knapp, Roser Vento-Tormo, Geoffrey Schiebinger, Peter W. Zandstra

**Affiliations:** School of Biomedical Engineering, University of British Columbia, Vancouver, British Columbia, Canada; Paul Albrechtsen Research Institute, CancerCare Manitoba, Winnipeg, Manitoba, Canada; Department of Biochemistry and Medical Genetics, Rady Faculty of Health Sciences, University of Manitoba, Winnipeg, Manitoba, Canada; Department of Mathematics, University of British Columbia, Vancouver, British Columbia, Canada; Michael Smith Laboratories, University of British Columbia, Vancouver, British Columbia, Canada; Wellcome Sanger Institute, Wellcome Genome Campus, Hinxton, Cambridge, United Kingdom; Department of Mathematics and Statistics, University of Maine, Orono, Maine, USA; Institut de recherche en immunologie et en cancérologie and Département de pathologie et biologie cellulaire, Université de Montréal, Montreal, Québec, Canada; Life Sciences Institute, University of British Columbia, Vancouver, British Columbia, Canada

## Abstract

T cells develop from multi-potent hematopoietic progenitors in the thymus and provide adaptive protection against pathogens and cancer. However, the emergence of human T cell-competent blood progenitors, and their subsequent specification to the T lineage, has been challenging to capture in real time. Here, we leveraged a pluripotent stem cell differentiation system to understand the transcriptional dynamics and cell fate restriction events that underlie this critical developmental process. Time-resolved single cell RNA sequencing revealed that cell-cycle exit, downregulation of the multipotent hematopoietic program, and upregulation of >90 lineage-associated transcription factors all occur within a highly co-ordinated and narrow developmental window. Computational gene-regulatory network inference elucidated the transcriptional logic of T lineage specification, uncovering an important role for YBX1. We mapped the differentiation cell fate hierarchy using transcribed lineage barcoding and mathematical trajectory inference and discovered that mast and myeloid potential bifurcate from each other early in haematopoiesis, upstream of T lineage restriction. Collectively, our analyses provide a quantitative, time-resolved model of human T cell specification with relevance for regenerative medicine and developmental immunology.

## Introduction

T cells are key effectors of the adaptive immune system that protect us from infection. They are also being used to treat cancer in the clinic^1^ and have tremendous promise for treating infection, autoimmunity and cardiovascular disease^2–4^. Understanding the gene regulatory programs and cell fate decisions that govern T cell development will not only improve our fundamental understanding of this crucial developmental process, but will also guide efforts to differentiate T cells in vitro, unlocking an unlimited supply of T cells for therapy^5^.

During fetal development, hematopoietic stem and progenitor cells (HSPC) arise from a cell type known as hemogenic endothelium in the aorta-gonad-mesonephros region (AGM) ^6^. HSPC differentiate into T cells in the thymus^7^. Decades of research have explored the process of T cell maturation using a combination of mouse models^8^, primary human thymic tissue^9,10^ and in vitro T cell differentiation systems^11^. Recently, single cell RNA sequencing experiments have helped uncover some of the key gene expression programs that control T cell maturation^9,10^. Though our understanding of the maturation process is improving rapidly^8,10^, the transcriptional programs and cell fate decisions that control the early developmental step of human T lineage specification from HSPC are less clear^12^.

Complicating our exploration of this process, is the versatility of HSPC in generating other hematopoietic cell types beyond T cells. T lineage specification is a rare event compared to other steps in development^12^. A small number of HSPC seed the thymus before undergoing rapid expansion and differentiation^12^. Due to their limited numbers, these early thymic seeding progenitors represent a minimal fraction of the total cell population, which has led to their sparse representation in profiling experiments of the primary thymus tissue, leaving important gaps in our knowledge of the lineage commitment process^10,13^. Comparing the cells at the earliest stage of specification to the T lineage directly to cells that have recently adopted alternate hematopoietic fates could reveal the transcriptional programs that drive the fate-determination process.

Although sampling primary human thymic tissue has greatly informed our knowledge of T cell development, it has not been possible to sample the same primary tissues with fine temporal resolution. Consequently, we do not know whether human T lineage specification from HSPC is a gradual or rapid temporal process. Pseudotime ordering of cells sampled at fixed time points suggests that T lineage specification occurs through multiple discrete waves where loss of a multipotent program precedes expression of T lineage-specific transcription factors^12,14^. In vitro T cell differentiation cultures can be initiated and sampled iteratively, presenting the opportunity to test the proposed multi-wave model by studying the dynamics of fate restriction in real time.

A related limitation of primary tissue samples is that they are not amenable to genetic perturbation, limiting our knowledge of the transcription factor programs that actually promote T lineage specification in humans. Thus, while the gene regulatory networks (GRNs) governing T lineage specification prior to TCR recombination has been studied in impressive detail in mouse^8^, equivalent understanding in humans is relatively lacking^15,16^. Beyond the well-established role for notch signalling, researchers have only validated important roles for a limited number of transcription factors such as GATA3 and TCF7^8^, suggesting the possibility for other contributory genes that are yet to be discovered.

Lastly, the point at which cells within the hematopoietic hierarchy become restricted to specific downstream lineages remains uncertain. Multipotent HSPCs are transcriptionally heterogenous and recent work suggests that downstream fate biases may already exist before cell surface markers are able to distinguish discrete cell types^17,18^. Moreover, recent work has shown that most combinations of lineage can branch directly from multipotent cells ^19,20^, blurring the strict bifurcating tree as classically depicted. The exact details of early lineage priming events thus remain poorly explored, and this is particularly the case for T cells and mast cells that are relatively sparsely represented in the in vitro assays that have been used to measure such differentiation potentials^21^.

Because it is technically impractical to sample cells from the same clones undergoing lineage specification at multiple timepoints in primary human samples, and considering that these samples are not amenable to genetic perturbation, alternate technology platforms are needed to address the aforementioned gaps in our knowledge of T cell fate specification. We recently developed a defined in vitro differentiation platform that allows us to capture HSPC emergence and T lineage specification starting from pluripotent stem cells (PSCs)^22^. We have previously shown that the system faithfully recapitulates key T cell developmental milestones and produces a broad complement of hematopoietic cell types with strong transcriptional resemblance to the primary blood cells that arise during human fetal development^22^. Importantly, nascent HSPC produced in this protocol are transcriptionally similar to primary definitive hematopoietic stem cells from the Carnegie stage 14/15 aorta-gonad-mesonephros (AGM), the major site of HSC emergence in vivo^22^.

Here, we used this platform to perform a sequencing time course, sampling cells every 24 hours over the period where HSPC emerge and become T cell progenitors. We used computational modelling, genetic perturbations and transcribed lineage barcoding to dissect the gene expression dynamics and cell fate decisions that control T lineage specification from stem cells.

We observed a rapid transcriptional overhaul where downregulation of a core cell-cycle-associated program co-occurs with upregulation of nearly 100 transcription factors within a ∼48-hour temporal window. We constructed a gene regulatory network model of human T cell specification which we used to generate experimentally validated in silico predictions, including revealing an important and previously unknown role for the transcription factor YBX1.

Using quantitative barcoding experiments, we made the surprising observation that mast cells share limited ancestral overlap with other myeloid cell types and we demonstrated that T cells occupy an intermediate position between mast and myeloid cells on the developmental landscape. Finally, we implemented an improved mathematical framework for estimating cell fate probabilities from barcoding scRNA-sequencing data. This allowed us to look back in developmental time and identify the earliest-acting transcription factor programs associated with hematopoietic fate specification. Collectively, this work provides a detailed picture of the gene expression trajectories and clonal dynamics that govern HSPC differentiation and T lineage specification.

## Results

### Gene expression changes associated with HSPC emergence and fate specification

We performed time-resolved scRNA-sequencing analysis of HSPC emergence and T lineage specification from human PSCs (Fig. 1a). We used a differentiation protocol that our laboratory previously developed^22^. Briefly, PSCs are differentiated into mesoderm and subsequently hemogenic endothelial cells through stepwise exposure to recombinant proteins and small molecules^22–24^. Next, the hemogenic endothelial cells are cultured in a media that promotes endothelial-to-hematopoietic transition (EHT) in plates coated with the notch ligand DLL4 and the cell adhesion protein VCAM1 for 7 days. Next, the hematopoietic cells that emerge during EHT are passaged on to DLL4 and VCAM1 coated plates and cultured in a media that supports T cell progenitor differentiation for an additional 7 days. To mitigate batch-to-batch variation in cell capture and sequencing, we initiated 14 differentiations, beginning with the EHT stage, on consecutive days. On day 14, we labelled cells from each differentiation timepoint with hashing antibodies^25^ and pooled samples for single cell capture and sequencing (Fig. 1a).

**Figure 1:**
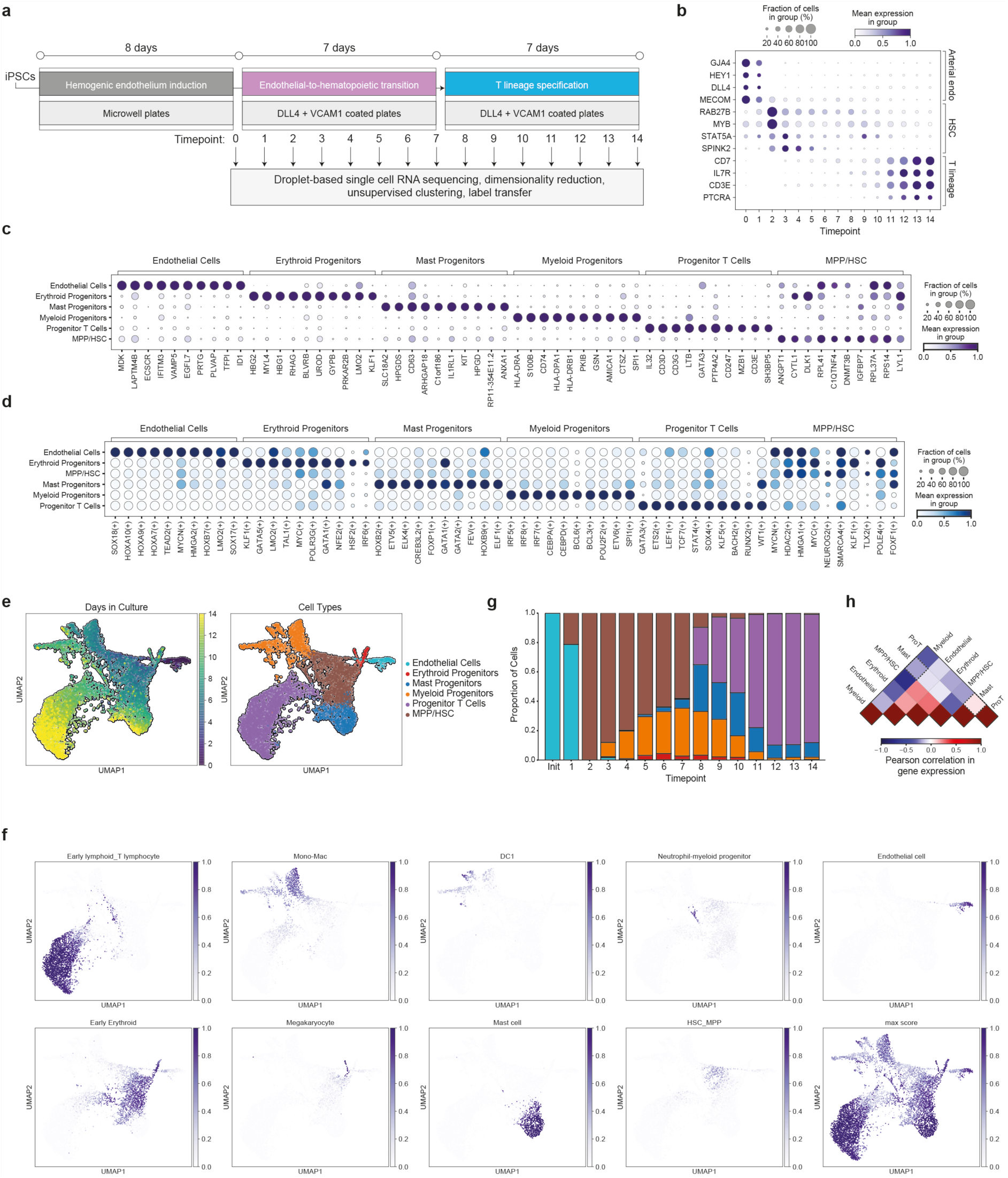
In vitro differentiation provides a transcriptionally accurate, temporally-resolved model of multi-lineage haematopoiesis. **a.)** Experimental design for scRNA-sequencing time course. **b.)** Temporal expression analysis of scRNA-sequencing data for marker genes associated with associated with key developmental milestones on the path to T lineage specification. **c,d.)** Differentially expressed genes (**c**) and differentially active SCENIC regulons (**d**) across unbiased leiden clusters used for cell type annotation. MPP/HSC = multipotent progenitors/ hematopoietic stem cells. **e.**) Projections of cells from scRNA-sequencing dataset annotated by differentiation timepoint (left) and cell type (right). Timepoint annotations are based on cell-hashing and reflect actual time in culture. **f.**) Projection of anchor-based integration scores for global transcriptional correlation with the specified primary cell type from a published primary fetal liver reference dataset. g.) Proportion of each cell type as a function of time point in the scRNA-sequencing dataset. **h.**) Pearson correlation in gene expression between cell types. The correlation between mast cell and progenitor T cell (Pro T) transcriptomes is higher than for mast cells and myeloid cells (indicated by dashed border).

After quality control and filtering^26^, we annotated cells by time point and analysed the temporal expression dynamics of genes associated with endothelial, hematopoietic, and T lineage identity (Fig. 1b). Mounting evidence suggests that HSCs arise from a precursor that transiently displays arterial endothelial identity^27,28^. We detected strong and broad expression of the arterial endothelium genes^27,28^ *GJA4, HEY1, DLL4* and *MECOM,* in the day 0 and day 1 samples (Fig. 1b). HSC genes^27,28^ *RAB27B, MYB, STAT5A* and *SPINK2* were sparsely expressed on days 0 and 1 and rapidly and strongly upregulated on day 2 before being gradually downregulated from day 3 onward (Fig. 1b). The T lineage genes *CD7* and *CD3E* were expressed at low levels in a small subset of cells as early as day 3 (Fig. 1b). Following passage into differentiation media that supports T cell development on day 7, the T cell genes^9,10^ *CD7, CD3E, IL7R* and *PTCRA* were all strongly and broadly upregulated, plateauing by day 13 (Fig. 1b). These expression dynamics indicate that cells in our culture system pass through an arterial endothelium state, transition through a HSPC intermediate and adopt a T lineage identity upon exposure to appropriate cytokines, faithfully copying key events observed in human development^10,27^.

To analyse this process at greater resolution, and identify additional cell types that are generated by our culture conditions, we performed unsupervised clustering and differential gene expression analysis (Fig. 1c-g). In addition to endothelial cells, HSPC and T cell progenitors, we also detected *HLA-II+* myeloid progenitors *HBG1/2+, LMO2+*, erythroid/megakaryocyte (Ery/Mk) progenitors and *KIT+, IL1RL1+, CD63+* mast cell progenitors^29^. This mast population is notable as the lineage hierarchy of human mast cell development has not been comprehensively characterized and the ancestral relationship between human mast cells, other myeloid cells and T cells is not known^21,29^.

In addition to differential gene expression analysis, we also inferred transcription factor (TF) activity in our dataset using SCENIC^30^ (Fig. 1d). Several of the TFs most strongly associated with each cell type in vitro have previously been identified in vivo^21,31–34^. Consistent with primary HE that can give rise to HSCs in human development^35^, endothelial cells in our data set showed strong activity of the late HOX genes *HOXA10*, *HOXA9* and *HOXA7* as well as *SOX17* (Fig. 1d). Ery/Mk progenitors had strong *KLF1, GATA5, LMO2* and *TAL1* activity (Fig. 1d). *IRF*, *C/EBP* and *BCL* family members were highly active in myeloid cells in addition to *SPI1*(Fig. 1d). *GATA1, GATA2, FEV* and *ETV5* were highly active in the mast lineage and T cell progenitors showed strong *TCF7, GATA3* and *ETS2* activity, all consistent with previous reports^21,34,36^(Fig. 1d).

We used an anchor-based integration strategy^37^ to perform an unbiased, transcriptome-wide comparison between our in vitro generated hematopoietic cells and a previously reported dataset from the human fetal liver^32^. This comparison corroborated our cell type annotations and confirmed that the cells generated in our protocol show strong correspondence to primary fetal blood cells (Fig. 1f). We verified this finding using a complementary logistic regression-based integration method (Fig. S1)

Next, to track the emergence of each population in real time, we analysed the relative abundance of each cell type at each time point (Fig. 1g, Fig. S1). Cells underwent a rapid transition from endothelial to MPP/HSC and myeloid progenitors started to emerge as early as culture day 3 and continued to accumulate, reaching 33% of the total culture by their peak at day 7(Fig. 1g). A small Ery/Mk population appeared transiently between culture days 5 to 10 (Fig. 1g). Mast cells began to arise on culture day 5 but rapidly increased in frequency following the passage from EHT media to T cell differentiation media after day 7 (Fig. 1g). T cell progenitors did not arise in appreciable numbers until after the transition to T cell differentiation media, but accounted for the majority of the culture by day 11 (Fig. 1g). This finding is consistent with T cell progenitors arising from the pre-existing HSC/MPP, rather than arising directly from HE.

Interestingly, the myeloid progenitor population rapidly declined following the passage, suggesting they cannot survive in the T cell differentiation media (Fig. 1g). In contrast, mast cell progenitors were able to persist until the final time point (Fig. 1g). We also compared global transcriptional identity between cell types^37^ and observed a strong correlation between the mast and T cell progenitor populations (Fig. 1h).

These analyses revealed population-wide temporal dynamics in gene expression and cell type abundances during in vitro differentiation. We next turned our attention to mapping the gene expression dynamics that promote cell fate specification within the annotated T cell progenitor population. We performed additional in-depth benchmarking of this cluster using well established stage-specific marker genes^38^ and by integration with three different single-cell thymus reference atlases^9,10,39^, using two different integration methods (Fig. S2). Strong upregulation of T cell specification and commitment genes^38^ including *BCL11B, PTCRA,* and *GATA3* and the concurrent downregulation of the multipotency associated genes *MEIS1, LYL1* and *BCL11A* alongside downregulation of the myeloid/multipotency associated gene *SPI1*, collectively confirm the progenitor T cell identity of this cluster (Fig. S2). Timing of these expression changes suggest that T lineage commitment occurs in the majority of these cells at approximately day 11 (Fig. S2). This is consistent with reduced expression of cell cycle genes which also begins around day 11(Fig. S2). Exit from cell cycle is a known hallmark of commitment^10,38^. Of note, culture media was replenished during the T cell differentiation period, supporting that cycle exit is not simply a consequence of depleted media.

When we examined transcriptome-wide cell type prediction scores in our PSC-derived T cell progenitor cluster, we observed high scores for primary T cell progenitor subsets and low or zero scores for myeloid subsets, verifying that this cluster predominantly comprises T cell progenitors (Fig. S2). While the highest prediction scores at each time point were overwhelmingly T-lineage, a small subset of cells also displayed modest Natural Killer cell prediction scores (Fig. S2).

Based on transcriptome-wide prediction scores from Lavaert et al^9^, the reference dataset with the highest representation of double-negative T cell progenitors, we observed a temporal transition in our PSC-derived progenitor T cell cluster where an initial population of early thymic progenitors (ETPs) gave rise to a committed T cell progenitor population which peaked at differentiation day 11 before the emergence of re-arranging progenitors (Fig. S2). Collectively, these data support the fidelity of in vitro differentiation platform for studying the early stages of human T cell development and suggest that the majority of cells within our annotated progenitor T cell cluster are committed to the T lineage at approximately day 11 of in vitro differentiation.

### Coordinated changes in Transcription Factor activity occur during a narrow temporal window in T lineage specification

Recent studies have examined the gene expression changes that occur during T cell development using primary human thymus samples^9,10,12,13^. Asynchronously differentiating cells were ordered in pseudo-time in order to infer developmental dynamics^9,10,12,13^. This approach allowed researchers to take genes known to be associated with a given lineage and map their expression levels to different stages of T cell maturation, but does not provide information on their expression dynamics in “real time” ^9,10,12,13^. Using our in vitro differentiation platform, we synchronized the onset of development and sampled cells every 24 hours for analysis. This approach offers temporal precision that is not available in previous analyses.

To capture the T lineage specification process, we focused on cells in the T cell progenitor cluster from days 8 to 14 of differentiation. We established an automated pipeline to identify dynamically changing transcription factor programs in an unbiased manner (Fig. 2a). First, we scored transcription factor activity in single cells using the SCENIC workflow^30^, which accounts for co-expression and the presence of binding motifs between TFs and their downstream targets. Next, we calculated mutual information^40^ between TF activity scores and time to find TF programs (SCENIC regulons) that vary temporally during T lineage specification. We then filtered TFs whose dynamics were well fit by a hill model (R^2^ > X, Fig. 2b). This pipeline returned 102 TFs that are strongly regulated during T cell development (Fig. 2c, Fig. S3). Most of these dynamic TFs (92/102) were upregulated over time (Fig. 2c). We extracted the K value from the hill fits for each dynamically regulated TF where K is the time at which each TF reaches the half-waypoint between minimal and maximal activity. These K values were unimodally distributed over a narrow time window, suggesting that transcriptional specification to the T lineage in humans is rapid rather than gradual. Ninety-five percent of the dynamically regulated TFs switched on or off between hour 252 and 296 of differentiation (Fig. 2d). Notably, this transcriptional overhaul occurs more than 3 days after cells were transferred into T cell differentiation media.

**Figure 2:**
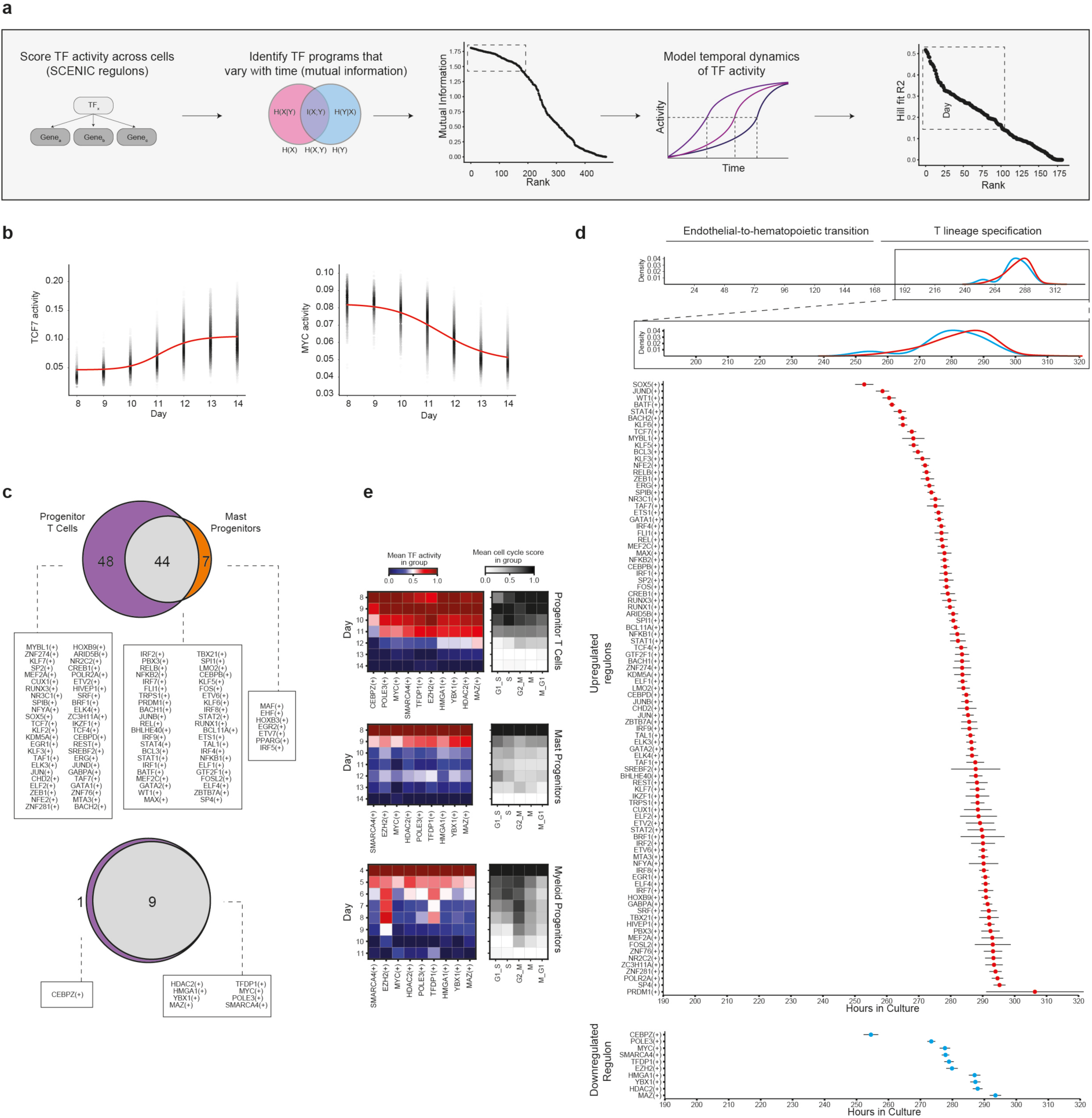
Widespread changes in transcription factor expression programs overlap with cell cycle exit during a narrow temporal window. **a.)** Computational workflow for identifying transcription factor (TF) programs with increasing or decreasing activity over time during T lineage specification. **b.)** Examples of a transcription factor programs identified by the computational pipeline in (a) that increase (left) or decrease (right). **c.)** List of transcription factors passing filter that increase (top) or decrease (bottom) during T cell differentiation (purple), mast cell differentiation (orange), or both (grey). **d.)** Plot of the k values (time where activity reaches half maximal) from hill fits for TFs that increase (top, red) or decrease (bottom, blue) during T cell differentiation. Error bars reflect standard deviation of k values. Histograms overlaid above the graphs show the density of K values over time. **e.)** TF activity plotted by day for TFs that decrease over differentiation time within each specified lineage. Corresponding transcriptional scores for cell cycle activity genes are also plotted.

We also performed this analysis during mast lineage specification (Fig. S3). Interestingly, the automated pipeline revealed that a highly overlapping set of TFs (9/10) are downregulated during T lineage and mast lineage specification (Fig. 2e). Although we were unable to recover TFs that were dynamically regulated during myeloid differentiation using the automated pipeline (the kinetics of myeloid differentiation were more heterogenous), when we manually inspected the 9 TFs downregulated during mast and T cell specification, we confirmed that all of them (9/9) were also downregulated during myeloid differentiation (Fig. 2e). These data show that a common set of TFs are downregulated in a coordinated time interval during fate specification from multipotent hematopoietic progenitors (Fig 2e). This coordinated program consists of *SMARCA4, EZH2, MYC, HDAC2, POLE3, TFDP1, HMGA1, YBX1* and *MAZ* (Fig. 2e). This gene set is enriched for genes associated with chromatin remodelling (6/9 genes, FDP 2.01 x10^-5,^ PANTHER overrepresentation test). Three of these genes (EZH2, TFDP1 and MYC) are involved in several GO biological processes related to cell cycle ^41^. We plotted transcriptional cell cycle activity over time for each lineage and found that expression of the genes in the core downregulated program was highly correlated with cell cycle activity temporally (Fig. 2e). As the downregulation of the common program coincides temporally with upregulation of lineage-associated TF programs, exit from multipotency, exit from the cell cycle, transcriptional regulation of chromatin remodelling genes, and specification to a downstream lineage appear to occur simultaneously during hematopoietic development in a narrow time window, contrasting the distinct waves that occur during early murine T cell development^14^. In addition to revealing this common TF program associated with an exit from multi-potent state, our analysis also provides a detailed resource of TF activity dynamics during T cell development. By fitting data to a hill model, we are able to estimate K values (the time that TFs reach half maximal activity) with an average precision of 2.44 hours (mean standard deviation of K).

Together, this unbiased analysis platform allowed us to describe the dynamics of individual transcription factor programs during T lineage specification with fine temporal resolution. This motivated us to next move from this descriptive dataset to a network-level understanding of the transcriptional programs that govern the specification.

### Gene regulatory network inference exposes an important role for YBX1 in T lineage specification

During cellular development, gene regulatory networks (GRNs) govern cell fate decisions^42^. Accurate GRN models can be used to predict the impact of perturbations in silico providing^43^ an accelerated avenue to discovering genes with a causative role in driving differentiation. We recently established a computational platform called IQCELL that can reconstruct GRNs from scRNA-seq data with high fidelity^44^. In brief, IQCELL selects active and dynamic regulons using SCENIC^30^, corrects for gene dropouts with MAGIC ^45^, scores candidate gene-gene interactions on the basis of mutual information^40^, and further filters interactions using a gene hierarchy based on the temporal order in which gene expression levels change during development (Fig. 3a). IQCELL utilizes the satisfiability modulo theory engine (Z3) to find Boolean logic functions that can explain the experimentally observed gene expression dynamics^44^. This results in an executable GRN capable of simulating cellular development using in silico models of wild-type and knock-out cells^44^.

**Figure 3:**
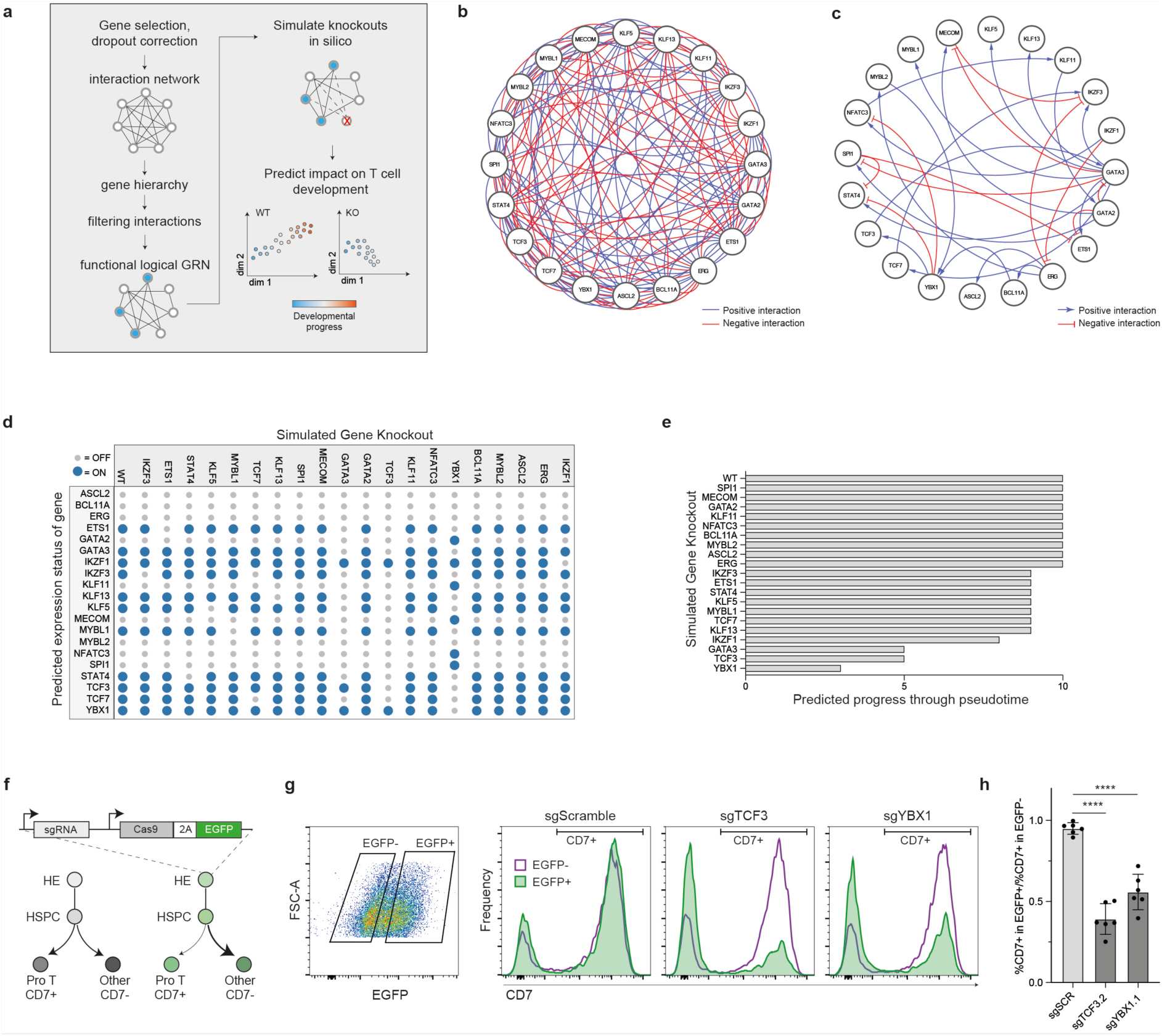
Gene-Regulatory Network Modelling reveals a role for YBX1 in T lineage specification from human pluripotent stem cells. **a.)** Schematic of the IQCELL platform for building executable gene-regulatory network (GRN) models from time-course scRNA-seq data. WT= wild type, KO= knockout. Graphs are illustrative only. **b.)** Set of all possible transcription factor interactions filtered by mutual information and by interaction hierarchy. **c.)** GRN inferred by IQCELL. GRN is inferred by constraining the interaction network in (b) to a logical, executable network that recovers temporal expression dynamics from the scRNA-seq dataset and that maximises mutual information between TF pairs. **d.)** In silico prediction of gene expression states following simulated gene knockouts. For each knockout, the predicted expression state of all TFs in the network once the system reaches a steady-state attractor is shown. **e.)** Predicted impact of TF knockouts on developmental progression through pseudotime, with a value of 10 representing measured progress for the unperturbed GRN. Pseudotime progression was divided into 10 bins, and the bin for the predicted steady-state attractor is plotted for each simulated knockout. **f.)** Experimental design for functionally validating the impact of TF knockouts predicted by IQCELL in PSC-derived cells. HE= hemogenic endothelium, HSPC = hematopoietic stem and progenitor cells, pro T= T cell progenitor. **g.)** Representative flow cytometry analysis at the end of the T cell progenitor differentiation culture phase. **h.)** Quantification of the perturbation experiment outlined in (f,g). Error bars reflect standard deviation of 6 independent transduction replicates, **** = adjusted P < 0.0001, Dunnett’s multiple comparisons test.

Here we applied IQCELL to our scRNA-seq data to model the GRN that drives T cell development from definitive, AGM-like, HSPCs (Fig. 3 a-c, Fig. S4). IQCELL predicted a GRN model where *YBX1* activates *TCF3*, which in turn activates *GATA3* which is a central hub in the network with multiple outgoing inhibitory and activating interactions (Fig. 3c).

In addition to inferring GRN structure, IQCELL can also simulate GRN dynamics over developmental time for wildtype and perturbed cells. We simulated the impact of knocking out each TF in the GRN in silico (Fig. 3d,e). This analysis predicts the impact of each knockout on developmental progress (Fig. 3e). The model predicted that the majority of single gene knockouts would have no or minimal impact on progression towards the T lineage. In contrast, IQCELL predicted that knocking out any of GATA3, TCF3 or YBX1 would markedly impede early T cell development (Fig. 3e). GATA3 knockdown has previously been shown to suppress human T lineage commitment^46^. TCF3 knockout impairs T cell differentiation at multiple stages in mouse thymocytes ^47,48^. A role for YBX1 in T cell development has not been demonstrated to our knowledge.

To validate key findings from our predicted GRN and to determine the impact of TCF3 and YBX1 perturbation on human T cell development, we established a polyclonal CRISPR-knock out workflow (Fig. 3f). 48 hours into the EHT culture phase, we delivered Cas9, an sgRNA and an EGFP reporter using an all-in-one lentiviral vector^49^. A positive control sgRNA targeting CD7 reduced the percentage of CD7+ cells by 16.1-fold compared to a non-targeting scrambled negative control sgRNA (Fig. S4). sgRNAs targeting NOTCH1 and IL7R, genes known to be essential for early T cell development^50,51^, reduced CD7+ T-lymphoid progenitor output by 6.1-fold and 2.6-fold respectively to the scrambled negative control sgRNA (Fig. S4). After validating the polyclonal knockout workflow using these controls, we tested 3 sgRNAs each targeting TCF3 and YBX1. We observed a significant reduction in T cell progenitor output for all sgRNAs (Fig. 3g,h, Fig. S4). Compared to the nontargeting control, the most potent YBX1 sgRNA decreased CD7+ T-lymphoid progenitor output by 56%, or 2.4-fold (P_adjusted_< 0.0001, Dunnet’s multiple comparisons test, Fig. 3g,h). The most potent TCF3 sgRNA decreased CD7+ T-lymphoid progenitor frequencies by 39%, or 1.7-fold (P_adjusted_< 0.0001, Dunnet’s multiple comparisons test, Fig. 3g,h). We verified that transducing the most potent YBX1 sgRNA into primary umbilical cord blood-derived HSPC also significantly reduces CD7+ T-lymphoid output in vitro (Fig. S4). In addition to decreasing T cell progenitor differentiation, the YBX1 sgRNA also increased the relative frequency of mast and myeloid cell output compared to a non-targeting control sgRNA (Fig. S4).

These data highlight the utility of the GRN inferred by IQCELL. Not only were we able to verify an important role for TCF3 in human T cell specification, we were also able to uncover a previously unreported function for YBX1 in T cell development. A role for YBX1 is not readily apparent from differential expression data, highlighting the ability of our executable logical GRN model to identify latent transcriptional regulators of differentiation.

### Lineage barcoding reveals that mast cell clonal ancestry overlaps more strongly with T cells than with other myeloid cell types

In the analyses described thus far, we leveraged our time-resolved scRNA-sequencing data to understanding gene expression dynamics in cells that we confidently assigned to a downstream hematopoietic fate via unsupervised clustering. But what about cells from early time points in our differentiation culture that exist within the diffuse HSC/MPP population? Transcribed lineage barcoding in mice revealed lineage biases within the HSC/MPP population but did not explore T lineage priming^18^. Another study using a distinct human PSC-to-T lineage differentiation protocol reported that T lineage-primed progenitors can arise directly from HE^52^, but it is not clear whether this is a default program, or reflects a unique property of the first wave of fetal T cell differentiation^13^. In contrast to this previous report, we did not detect a clear progenitor T cell population early during EHT (Fig. 1g), but fate biases could already exist within the HSC/MPP cluster. We therefore explored how and when fate restriction occurs during earliest post-EHT developmental time points.

To understand how downstream cell fate outcomes are shaped within the upstream HSC/MPP population, we performed a transcribed lineage barcoding experiment. We adapted the recently pioneered Lineage And RNA RecoverY (LARRY) method ^17^, creating a new lentiviral barcoding pool that is robustly expressed in human PSC-derived hematopoietic cells. Our construct library comprises a bi-directional promotor driving expression of two transcripts; i.) a transmembrane reporter (tNGFR), and ii.) an EGFP mRNA that contains a 16-mer randomized barcode upstream of the poly A tail (Fig. 4a).

**Figure 4:**
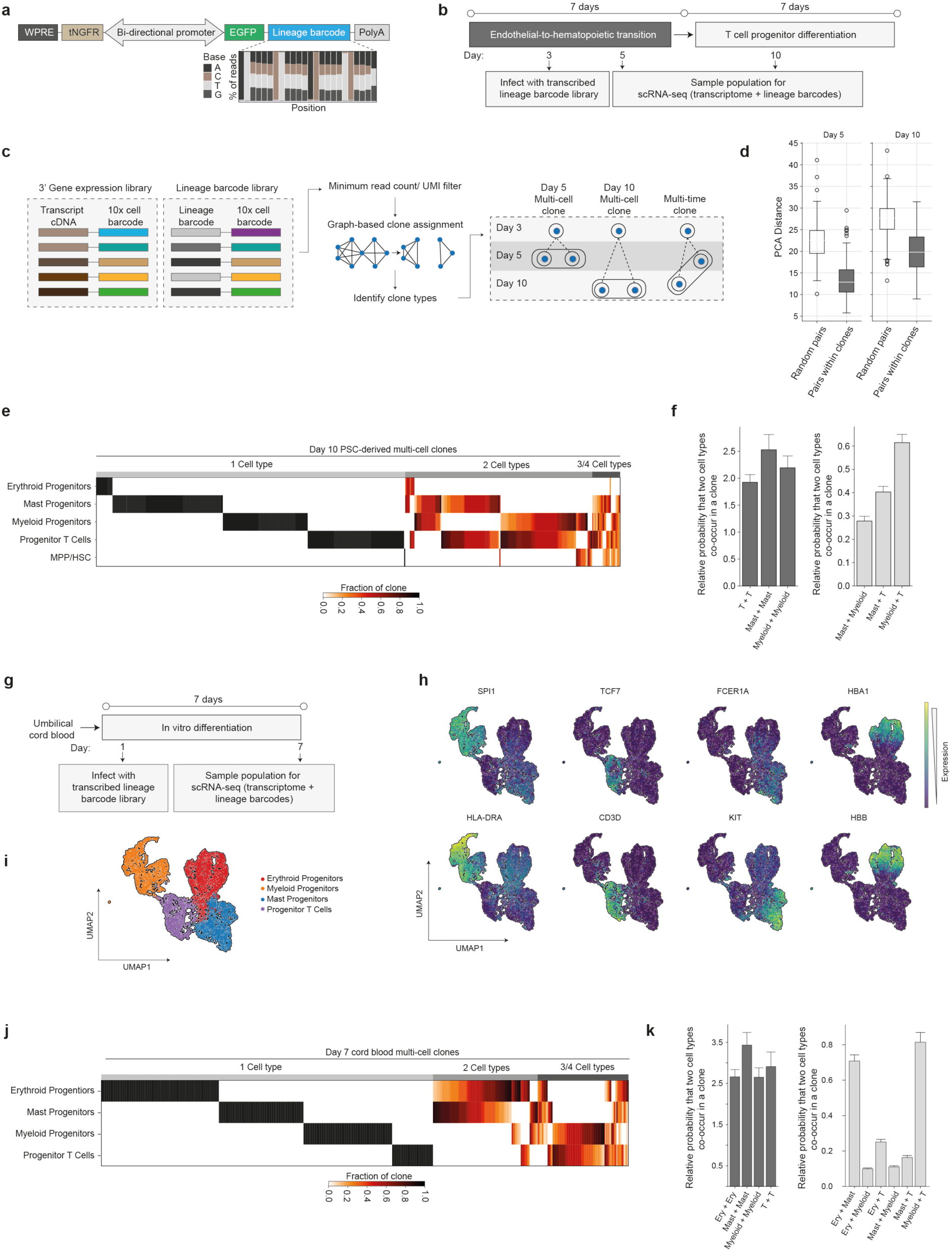
Transcribed lineage barcoding demonstrates that mast progenitors share more clonal overlap with T cells than with myeloid cells. **a.)** Design of transcribed lineage barcoding vector library, including high-throughput sequencing verification of barcode library diversity. Because the barcodes are proximal to the poly-A tail, their sequences can be recovered using a standard 10x Genomics cell capture. **b.)** Experimental workflow for transducing the barcoding library into PSC-derived cells during endothelial-to-hematopoietic transition culture. **c.)** Overview of analysis workflow for assigning clonal identity and transcriptomes to single cells. **d.)** Quantification of global transcriptional similarity between pairs of cells in day-5 and day-10 multi-cell clones from PSC-derived, in vitro differentiated cells. To create a reference control dataset, the same number of cell pairs that were present in true clones were randomly assigned to simulated control clones. **e.)** Distribution of cell types within day-10 multi-cell clones from PSC-derived in vitro differentiations. Vertical columns represent individual clones. **f.)** To quantify enrichment of cell-type combinations within day 10 multi-cell clones from PSC-derived differentiations, we measured the co-occurrence of cell type combinations within real-clones and normalized these values to the frequency of co-occurrence of cell types within simulated clones generated by randomly assigning cells to clones of the same size and number as the true-clone dataset. We performed this analysis for pairs of the same type (left) or different types (right) for mast, myeloid, and pro T cells where we had sufficient numbers for meaningful quantification. **g.)** Experimental workflow for transcribed lineage barcoding study conducted on CD34-enriched umbilical cord blood. **h.)** Expression of representative markers genes for myeloid (SPI1, HLA-DRA), T cell progenitor (TCF7, CD3D), mast cell (FCER1A, KIT) and erythroid (HBA1, HBB) following in vitro differentiation of umbilical cord blood for 7 days in multi-lineage permissive conditions. **i.)** Cell type annotations for umbilical cord blood cells differentiated for 7 days in vitro. **j.)** Distribution of cell types within day-7 multi-cell clones from umbilical cord blood-derived in vitro differentiations. Vertical columns represent individual clones. **k.)** Quantification of shared clonal ancestry amongst umbilical cord blood-derived cells after 7 days of in vitro differentiation using the same analysis method as (f).

We transduced differentiating PSC-derived cells on day 3 of the EHT culture phase. On day 5, we harvested half of the non-adherent cells for scRNA-seq (day 5 sample) and allowed the remaining cells to continue differentiation. After passaging the culture into T cell differentiation conditions on day 7, we sampled the remaining cells for a second scRNA-seq time point at day 10 (day 10 sample, Fig. 4b). We generated a 3’ gene expression library using the standard 10x genomics protocol and created a second targeted sequencing library using a primer pair that specifically amplifies barcoded transcripts (Fig. 4c). To correct for PCR artefacts and robustly assign cells to their respective clones, we filtered barcodes by read and Unique Molecular Identifier (UMI) counts and established a graph-based clone assignment pipeline (Fig. 4c, Fig. S5).

Following clone assignment, we identified 1389 total multi-cell clones including 850 day 5 multi cell clones, 451 day 10 multicell clones and 277 multi-time clones (clones containing at least one cell each from day 5 and day 10). Pairs of cells within clones had substantially more similar transcriptomes compared to randomly selected cell pairs confirming that our barcoding strategy is able to recover the correlation between transcriptional state and clonal identity (Fig. 4d).

Lineage barcoding gives us the ability to quantify ancestral overlap between cells fated towards the T, mast and myeloid lineages. Beyond T-cells, mast cells were of particular interest in our dataset. Mast cells contribute to vasodilation, angiogenesis and pathogen defence but also play a key role in allergic responses and asthma^53^. However, because these cells are mainly tissue-resident, they are poorly represented in peripheral blood samples and their ontogeny remains poorly understood^29,53–57^. Their high abundance in our in vitro differentiation cultures affords the opportunity to resolve their position in the lineage hierarchy and to explore the gene expression programs that promote their specification from HSPC as the expense of alternate fates.

While mast cells have traditionally been classified as a subset of the myeloid branch^29,53–56^, a recent study in mouse suggests that mast fate diverges from neutrophil/monocyte/macrophage fate earlier in haematopoiesis than previously appreciated^21^. This early specification was strongly associated with GATA1 expression^21^. GATA1+ progenitors could give rise to mast cells and eosinophils but lacked monocyte, macrophage, neutrophil and lymphoid potential, suggesting mast and T lineage potential may also segregate early during murine haematopoiesis^21^. In a separate study, enforced GATA3 overexpression diverted mouse T cell progenitors to the mast lineage, suggesting relatively high plasticity between these two cell types^57^. Our own transcriptional profiling of human hematopoietic progenitors revealed stronger transcriptional correlation between mast and T cell progenitors compared to mast progenitors and other myeloid cell types (Fig. 1h). To move beyond this correlational data, we leveraged our barcoding dataset to quantify the true ancestral overlap between cells of each lineage.

We examined the distribution of cell types within individual day 10 multi-cell clones (Fig. 4e). A substantial proportion of clones containing pro T cells also contained myeloid cells, verifying that pro T cells arise from multi-potent ancestors (Fig. 4e). Similarly, many of the clones with T lineage potential also gave rise to mast cell progenitors. Interestingly, we observed less clonal overlap between mast and myeloid (neutrophil, monocyte, macrophage, dendritic cell) lineages (Fig. 4e). To formally quantify shared ancestry between lineages, we calculated the distribution of cell fate combinations within pairs of cells in true clones and normalized this to the distribution of cell fate combinations in pairs of cells chosen at random and assigned to simulated clones (Fig. 4f). True clones contained pairs of cells with the same fate (e.g. T cell, T cell pairs) at a substantially higher frequency than expected by chance alone (Fig. 4f). When we examined pairs of cells from different lineages, we found that Myeloid/T and Mast/T pairs were significantly more enriched than Mast/Myeloid pairs (Fig. 4f). These data demonstrate, that in our in vitro differentiation system, mast cells share more overlapping clonal ancestry with T cells than they do with other myeloid cell types. They also demonstrate that myeloid cells share more ancestral overlap with T cells than they do with mast cells (Fig. 4f). Together, these data position mast cells as at the opposite end of the hematopoietic fate spectrum from other myeloid cells.

To determine whether the lack of clonal overlap between mast and myeloid cells is PSC-specific, or generalizable to post-natal haematopoiesis, we performed a second transcribed lineage barcoding study using umbilical cord blood (UCB) (Fig. 4g). After 7 days of in vitro differentiation, we detected myeloid, mast, erythroid and T cell progenitors by scRNA-sequencing (Fig. 4h,I). Consistent with PSC-derived cells, UCB-derived T cell progenitors occupied an intermediate position in between mast and myeloid cells in gene expression space (Fig. 4I).

When we examined the distribution of cell types within UCB-derived day 7 multi-cell clones, we observed many clones comprising both erythroid and mast cells or both myeloid and T cells, but a dearth of clones containing both mast and myeloid cells together (Fig. 4j). When we quantified ancestral overlap by normalizing the co-occurrence of fates in true clones to the co-occurrence of fates in cells randomly assigned to clones, we determined that erythroid/mast pairs and myeloid/T cell pairs were both >5 times more likely to occur than myeloid/mast pairs (Fig. 4k, P <0.0001, Mann Whitney U test). This UCB lineage tracing experiment strongly supports our observation in PSC-derived cells that the mast and myeloid cell fate trajectories lie at opposite poles of the hematopoietic differentiation continuum. These findings also corroborate evidence from murine haematopoiesis that mast and erythroid cells are clonally adjacent cell types^21^.

Although paired transcriptional profiles and lineage barcoding data are not accessible for the primary human thymus, we examined gene expression correlations in a published single cell thymus atlas^10^ that contains T cell progenitors as well as mast cells, megakaryocytes, macrophages, monocytes, neutrophil-myeloid progenitors, dendritic cells and erythrocytes. Pearson correlation analysis revealed that mast cells were more similar to early T cell precursors (ETPs) than any other hematopoietic cell type in the reference dataset ^10^ (Fig. S5), supporting the conclusions of our in vitro lineage barcoding analysis.

### Mast and myeloid potential segregate from each other early in hematopoietic development within a population that retains T lineage potential

During hematopoietic differentiation from PSCs, we determined that cell fate specification was already underway during the EHT differentiation phase (Fig. 4f). We therefore asked whether cells might already display gene expression differences at day 5 of differentiation that are associated with the downstream fates we observed at day 10. To answer this question, we used our lineage barcoding information to explore the transcriptional profiles of day 5 cells in multi-time clones (Fig. 5a-e). The vast majority of day 5 ancestors of every downstream day 10 cell type were MPP/HSCs (Fig. 5b). We then annotated day 5 ancestors by their day 10 fates (Fig. 5c). Day 5 ancestors fated to give rise exclusively to T cells and day 5 ancestors fated to give rise to multiple cell types were distributed diffusely in gene expression space throughout the MPP/HSC population (Fig. 5c). In contrast, we observed a striking divergence between day 5 ancestors destined for mast vs myeloid fates (Fig. 5c). Erythroid-fated cells were already strongly segregated by day 5 (Fig. 5c). Interestingly, the small number of cells fated to retain their HSC/MPP identity at day 10 occupied a narrow cleft between the mast and myeloid fated cells and may represent a rare cell state capable of self-renewal (Fig. 5c).

**Figure 5:**
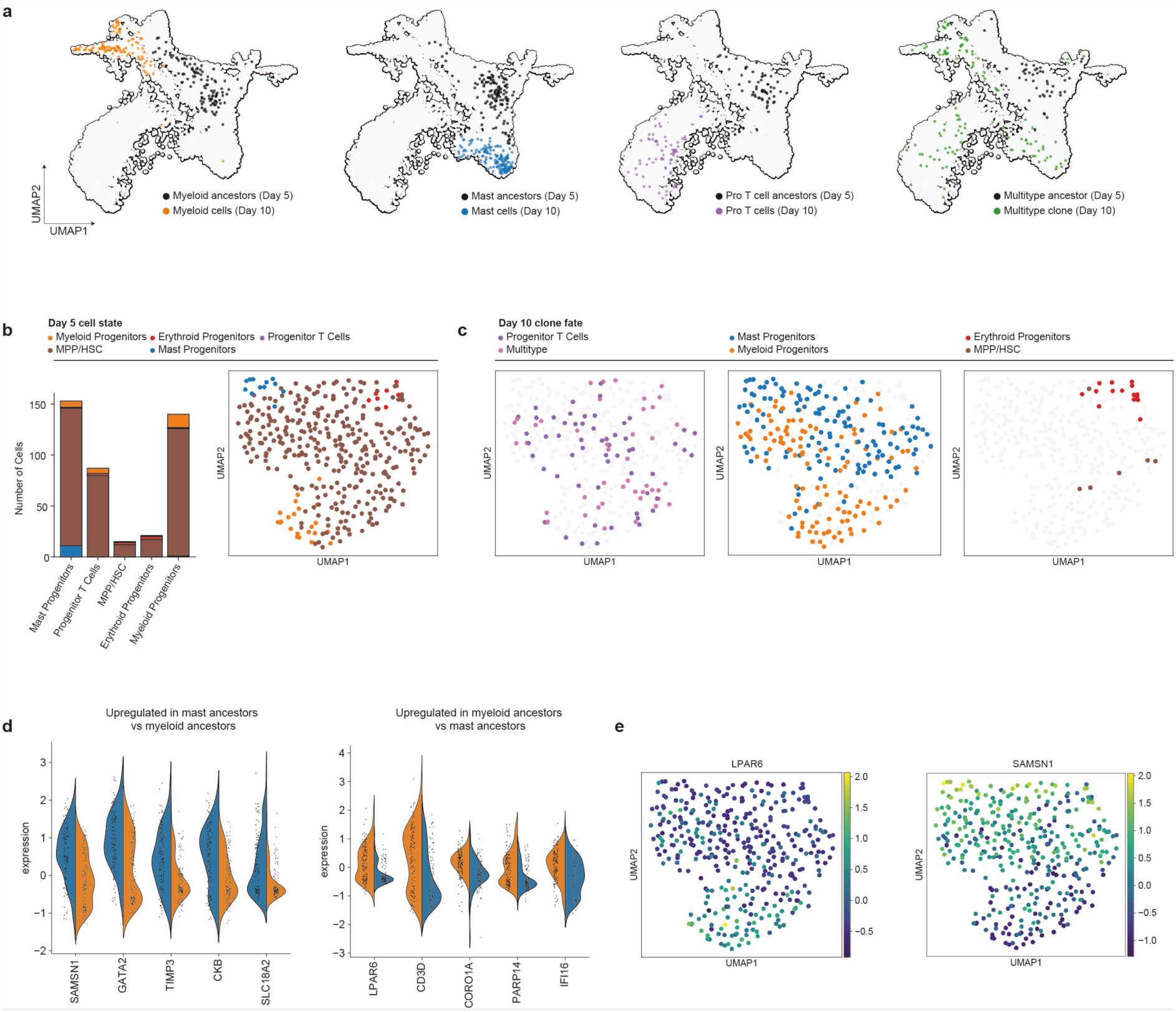
Mast and Myeloid potential are already segregated within the T-competent MPP/HSC pool at day 5. **a.)** Day 5 ancestors and day 10 progeny for multi-time clones. Barcoded cells were regressed to the scRNA-sequencing time-course dataset for visualization. **b.)** Visualization of the transcriptional state of day 5 cells in multi-time clones. Bar graph (left) depicts the day 5 state (colour) of cells based on their day 10 fate (column). UMAP (left) is a new projection generated using only the day 5-cells belonging to multi-time clones. Cells are annotated by their day 5 identity. **c.)** Projections from (b) annotated based on the day 10 fate of cells in the clone. **d.)** Top 5 differentially expressed genes that are enriched in the day 5 ancestors of day 10 mast cells compared to day 10 myeloid cells (left) or enriched in the day 5 ancestors of day 10 myeloid cells compared to day 10 mast cells (right). **e.)** Day 5 cells are coloured based on expression of the genes most strongly associated with downstream myeloid (left) or mast (right) fate.

Given the clear separation between mast-fated and myeloid-fated cells, we examined differentially expressed genes across the ancestors of each populations. On average, day 5 cells fated to the mast lineage expressed higher levels of *SAMSN1*, *GATA2*, *TIMP3, CKB and SLC18A2* whereas cells fated to other myeloid types expressed higher levels of *LPAR6, CD3D, CORO1A, PARP14* and *IFI6* (Fig. 5d,e). In mice, *Gata2* (and *Gata1*) expression marks specification towards mast, eosinophil, erythroid and megakaryocyte fates and loss of monocyte and neutrophil potential^21^ and SAMSN1 and SLC18A2 expression are highly enriched in mast cells (bioGPS mouse). LPAR6 is a lysophosphatidic acid (LPA) receptor. LPA receptor expression is high on human myeloid progenitors and LPA stimulation significantly increases myeloid differentiation from HSPCs^58^. In summary, we identified a segregation between mast-potent and macrophage/monocyte/DC/neutrophil-potent cells early in hematopoietic differentiation prior to the emergence of a distinct T-lineage-biased population. We report a set of genes that are already differentially expressed in the ancestors of the downstream cell types at least as early as day 5 in our in vitro differentiation system.

### Mapping the continuous landscape of fate specification with Lineage-informed Trajectory Inference

We next sought to explore hematopoietic fate specification from PSC-derived HSPC in greater detail. To extend our insights beyond the limited number of cells in multi-time clones and capture the temporal dynamics of fate restriction, we applied several lineage-resolved trajectory inference algorithms, including Lineage Optimal Transport (OT) ^59^ and CoSpar^60^. LineageOT is an extension of WaddingtonOT ^61^, which uses optimal transport to identify ancestor-descendant relationships in scRNA-seq time-courses. CoSpar extends multi-time clonal information to the rest of cells by propagating lineage information over a nearest-neighbourhood graph^60^. The two algorithms produced largely similar trajectory estimates, but there were some subtle differences (Fig. S6). In particular, LineageOT inferred that day 10 myeloid cells largely arose from pre-existing myeloid cells, whereas CoSpar inferred that they came from MPP/HSC (Fig. S6). CoSpar may be prone to sampling bias due to differences in cellular growth rates^62^. LineageOT does not use multi-time clone information directly (it only uses clonal relationships within time-points) and it relies on estimates of cell growth rates. Errors in growth rate estimates will hinder the performance of LineageOT.

We therefore utilized experimental measurements of cell growth rates (see methods), and used a modified version of LineageOT that incorporate multi-time data - a method called Multi-Time LOT (MT-LOT) (see methods and Bonham-Carter et al.^62^). The results from MT-LOT reflect a midpoint between the previous predictions from CoSpar and LOT and we proceeded using this experimentally-supported inference method. The fate trajectories predicted by MT-LOT allowed us to calculate the probability that each cell in our dataset is destined for a given fate (Fig. 6a).

**Figure 6:**
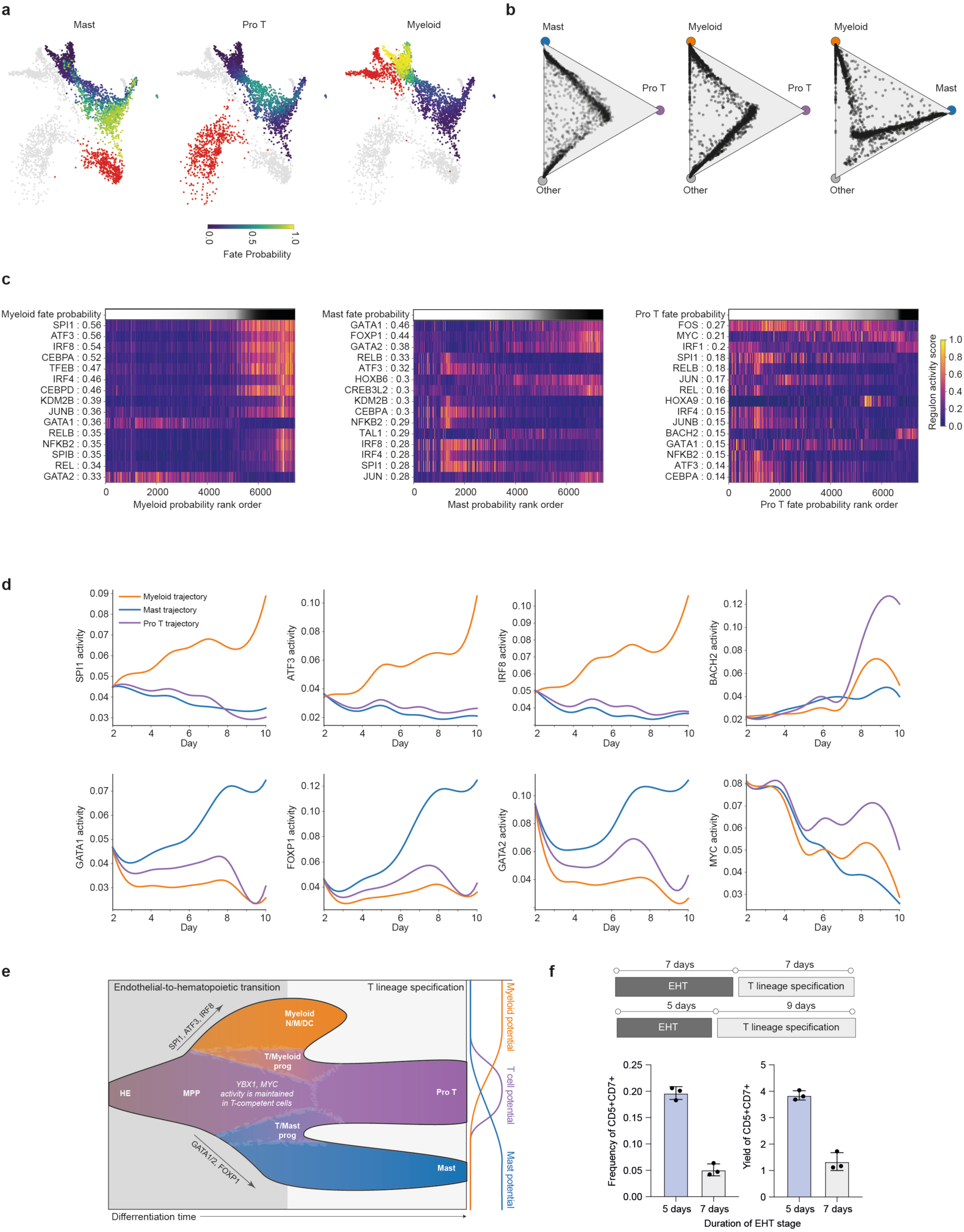
Mapping transcription factor programs associated with early specification to downstream hematopoietic lineages. **a.)** UMAP projections of day 5 cells from the lineage barcoding experiment coloured by the probability that they are fated to the specified downstream lineage at day 10 calculated using MT-LOT. Day 10 cells for the indicated lineage are coloured red. **b.)** Fate triangle plots demonstrate the probability that a given day 5 cell is fated for each other two lineages or other fates. **c.)** Transcription factor programs associated with Myeloid, mast and T cell fate probability. TFs were identified by calculating TF activity scores using SCENIC and calculating mutual information between SCENIC TF activity and MT-LOT fate probabilities. Numbers following the TF name indicate the mutual information score. Each column corresponds to a cell and cells were ordered from left-to-right based on increasing probability that the cell is fated for the specified lineage. **d.)** Activity of TFs most strongly associated with mast, myeloid or T cell fate are plotted over time by trajectory, as determined by MT-LOT. **e.)** Model of the fate specification during in vitro differentiation from PSC-derived hemogenic endothelial cells. The model is reflective of the clonal dynamics and gene expression programs observed in our study. **f.)** Experimental validation of the model prediction that shortening the EHT culture phase will accelerate differentiation of CD5+, CD7+ T cell progenitors. Bar graphs quantify flow cytometry analysis of progenitor T cell differentiation cultures carried out as indicated. Error bars are standard deviation of n=3 differentiation replicates.

MT-LOT analysis strongly supported our previous observation that cells with a high probability of giving rise to mast cells were clearly segregated from myeloid fated cells (Fig. 6a). Pro T cell ancestry was concentrated in a central region of the HSC/MPP population, and partially overlapped with the myeloid ancestors and mast ancestors (Fig. 6a). We plotted day 5 cells based on the relative probability that they will give rise to progeny for each of two lineages (Fig. 6b). We observed a front of cells occupying a continuum between pro T and mast fates and between pro T and myeloid fates at day 5 (Fig. 6b). In stark contrast, mast and myeloid fates are clearly separate branches in the fate probability space (Fig. 6b). This trajectory inference, together with the previously described clonal barcoding results (Fig. 4f), supports a revision to the classically-held view that mast cells arise from a common-myeloid progenitor that has already lost T-lineage potential. To the contrary, mast and myeloid fate potential can diverge even before T lineage specification occurs, and mast cells and T cells share more common ancestry than expected based on previous models^21^.

We next turned our attention to dissecting the transcription factors programs that are associated with mast vs myeloid fate decision making within the MPP. We combined the fate probability scores computed by MT-LOT with transcription factor activity scores calculated by SCENIC to identify early-acting transcription factors that are associated with lineage specification and plotted their temporal dynamics (Fig. 6c,d, Fig. S6). Unbiased mutual information analysis revealed a set of TFs with sharply opposing associations for mast and myeloid probability (Fig. 6c). *GATA1, GATA2* and *FOXP1* had strong positive associations with mast probability and strong negative associations with myeloid fate probability (Fig. 6c). *SPI1, IRF8, ATF3* and *CEBPD* were among a larger set of TFs that were positively associated with myeloid fate and negatively associated with mast fate (Fig. 6b). For many of these TFs, including ATF3, IRF8, GATA1, GATA2 and FOXP1, cells on the T lineage trajectory displayed an intermediate activity level in between the mast and myeloid trajectories (Fig. 6c). Only a small number of TFs, including *MYC* and *BACH2*, had positive associations with T cell fate (Fig. 6b), and in general TF associations with the T lineage were weaker than associations with mast or myeloid fate at this early time point (Fig. S6). *MYC* is part of the core program that we identified above that is downregulated during lineage specification from HSC/MPP. Of the few TFs were strongly associated with the T cell fate, their activity diverged later in differentiation time compared to TFs associated mast or myeloid specification (Fig. 6d).

Our lineage barcoding data and MT-LOT show that, in our differentiation system, HSPC rapidly become distributed across a developmental axis that divides mast and myeloid potential (Fig. 5c, Fig. 6 a,b). At intermediate positions along this continuum, cells lose mast or myeloid potential but retain their ability to become T cells whereas at the poles, cells become restricted to only mast or myeloid fate (Fig. 4f, Fig. 6a,b). This suggests that the earliest transcriptional events that restrict mast and myeloid fates from each other do not prohibit access to the T cell fate (Fig. 4f, Fig 6a,b,e). Progenitor T cells then rapidly emerge from following passage into pro T-permissive culture conditions after day 7 (Fig. 1g). This is an important revision to the previously proposed model that T lineage priming occurs upstream, within a subset of HE cells prior to undergoing EHT^13,52^. Our data instead support a scenario where, in the absence of specific T-lineage priming cues, MPP progressively lose T cell potential as they progress along a transcriptional continuum towards mast or myeloid fates. Importantly, we did not observe a discrete border dividing cells into distinct populations with pure fate outcomes, but rather a gradient of fate potential that correlates with the magnitude of transcription factor dosage (Fig. 5e, Fig. 6a).

Although our EHT conditions did not support substantial T lineage development prior to passaging cells into T cell differentiation media, they do allow commitment towards the mast and myeloid lineages by culture day 7 (Fig. 1g). Our revised model suggests that T cell potency is irreversibly lost in cells that have travelled to the extremes of the mast or myeloid continuum before the day 7 passage (Fig. 6e). This model therefore predicts that shortening the EHT culture phase from 7 days to 5 days should accelerate the production of pro T cells without compromising on overall yield. As predicted, reducing the duration of EHT significantly increased frequency of CD5+,CD7+ pro T output on day 14 (Fig. 6f), supporting the fact that progenitors in the central region of the mast vs myeloid commitment continuum retain their T cell potency and can undergo lineage specification in the appropriate signalling environment (Fig. 6c-e).

Collectively, our scRNA-sequence time-course, CRISPR knockout experiments, transcribed lineage barcoding studies and mathematical trajectory inference provide a detailed picture of how T lineage cells emerge from early definitive HSPCs, and how mast and myeloid fates become segregated early during haematopoiesis. These data have shed new light on the gene expression programs that drive T lineage specification and delineated their temporal dynamics with fine resolution.

## Discussion

In this investigation, we leveraged an in vitro PSC differentiation system as a tool for unravelling the gene expression programs and lineage hierarchy at the core of T cell specification. We note that this system does not replicate all aspects of the complex microenvironments where HSPC and T cells emerge^5^. However, this system presents substantial benefits, effectively capturing the minimum essential signals required for development, thereby enabling the study of lymphopoiesis in an isogenic and scalable context^22^. This controlled environment is highly conducive to regular sampling and experimental perturbations, offering enhanced temporal resolution of human T cell development. It also provides an unlimited source of cells at the earliest stage of lineage specification, allowing us to elucidate previously unknown drivers of this process.

In addition to utilizing a scalable and developmentally facile in vitro differentiation method, we also established a powerful new integrated experimental and analytical framework for identifying gene expression programs and clonal restriction events that underlie cell fate determination. This study provides first demonstrations that IQCELL^44^ and MT-LOT can each uncover previously unknown regulators of cell fate decision making. We also provide important improvements to the nascent field of transcribed lineage barcoding^17^. We established a barcoding vector library that is strongly and persistently expressed in human hematopoietic cells and created a robust computational analysis pipeline for assigning clonal IDs to single cells. We utilized a new metric to quantify shared ancestry between lineages, and we apply an updated mathematical trajectory inference method that leverages such lineage information. We anticipate these tools will have broad applicability for studying the gene expression programs and cell fate decisions that govern cellular differentiation in myriad developmental systems. In the case of T lineage specification, this integrated pipeline has already elucidated important developmental features that we discuss below.

Using this time-resolved transcriptional data and IQCELL, we discovered a previously unknown role for YBX1 in T cell development which we verified with perturbation experiments in PSC-derived and primary HSPCs. Because YBX1 is not strongly enriched in post-specification T cell progenitors, simple differential expression analysis would not likely have nominated this TF as a driver of T cell differentiation. In contrast, IQCELL predicted that YBX1 is a key node in the T cell specification GRN that acts up stream of TCF3, and in-turn, GATA3, a central hub in T cell development. We also observed that YBX1 expression is downregulated as cells leave the multipotency state and exit the cell cycle. Our data demonstrated that exit from the cycling, multipotent state during the early days of the EHT culture phase is associated with a loss of T cell potency as cells become committed to mast or myeloid fates. Knocking out YBX1 in differentiating cord blood cells reduced T cell output while causing a concomitant increase in mast and myeloid output. Considering these findings together, we propose that the cells which retain T cell potency for the duration of the EHT culture phase, rather than fully committing to alternate fates, are ones that persistently retain expression of YBX1 and remain in the multipotent, cycling state at least until the beginning of the progenitor T cell differentiation phase. Future work is required to determine whether persistent YBX1 expression is the result of specific microenvironmental cues, underlying epigenetic or transcriptional differences, or is a stochastic phenomenon.

Another unexpected finding in our analysis was the early divergence in mast potential from other myeloid cell fates. Mast cells have long been recognized for their central role in allergic reactions and pathogen defence, but their lineage trajectory from HSPC, and their clonal relationship to the T lineage has remained unclear^21,29,53–57^.

Previous sorting experiments in mouse showed that GATA1+ cells did not have T cell potential but could produce mast, basophils, erythroid and megakaryocyte cells ^21,57^. Here we show that, in our differentiation platform, T cells and macrophages/monocytes or DCs are more closely related on the clonal family tree than these myeloid cell types are to mast cells. Our data suggest that differentiation to the mast lineage proceeds, first through losing, macrophages, monocytes and DC potential, and then, upon reaching sufficiently elevated levels of GATA1/GATA2 activity, undergoing mast commitment and losing T cell potential. The improved understanding of mast cell development elucidated in our study could help us make them efficiently in vitro, offering a renewable source of mast cells for research and potentially for cell therapy.

Cellular differentiation depends on context. In vitro and in vivo, developing cells themselves contribute key paracrine, autocrine and juxtracrine signals to that context. For example, we found that in vitro T cell differentiation efficiency is highly dependent on cell seeding density^22^. Despite the importance of context, hematopoietic lineage potential is often investigated in vitro by isolating single cells and enumerating their downstream output. This approach inherently removes each clone from its relevant multicellular environment. In contrast, lineage barcoding allowed us to study cell fate outcomes without disrupting the multicellular microenvironment.

We leveraged PSCs as a scalable and isogenic tool to discover an unappreciated role for YBX1 in T cell development and to learn that myeloid cells and T cells share more overlapping clonal ancestry than myeloid cells and mast cells. We subsequently verified that these two findings hold true for primary HSPC. This affirms that high resolution profiling of PSC to T cell differentiation can be applied, not only towards improving the efficiency of cell therapy manufacturing, but also as a discovery engine to learn the principles governing primary haematopoiesis.

## Methods

### Pluripotent stem cell culture and differentiation

#### PSC culture

PSC differentiation experiments were performed using iPS11, a human iPSC line that was derived from foreskin fibroblasts using episomal reprogramming vectors. This line was obtained from Alstem Cell Advancements and grown as previously described. Cells were cultured on tissue-culture treated 6-well plates coated with Geltrex (Life Technologies, A1413302) at 37°C for 1 hour at 1mL per well. Cells were grown in mTeSR1 (STEMCELL Technologies, 85850) with penicillin-streptomycin (0.5% v/v, Invitrogen, 15140122).

Cells were fed daily with fresh complete media and were grown at 5% CO2, 37°C. To thaw frozen cells, mTeSR1 was supplemented with the ROCK inhibitor Y-27632 (STEMCELL Technologies, 72308, 7.5 μM final). Cells were partially dissociated to small aggregates/clumps for passaging with 1× TrypLE Express (Thermo Fisher Scientific, 12605028) at 37°C. Cells were incubated in TrypLE Express for 2 to 4 min and then gently scraped. For the 24 hours that followed passaging, mTeSR1 was supplemented with the ROCK inhibitor Y-27632 (STEMCELL Technologies, 72308, 5 μM final).

#### Hemogenic endothelium differentiation

To generate CD34+, hemogenic endothelial cells from iPS11, we followed our previously published protocol without modification. For more details and the composition of each media formulation, see Michaels et al 2022 ^22^. Prior to harvesting PSCs for differentiation, AggreWell 400 six-well plates (STEMCELL Technologies, 34425) were prepared by rinsing with AggreWell Rinsing Solution (STEMCELL Technologies, 07010).

PSCs were grown to reach high confluency (>90%) prior to initiating differentiations and dissociated to a single cell suspension using TrypLE. Following, dissociation and cell counting, cells were centrifuged at 200g for 5 min. Supernatatnt was aspirated and cell pellets were resuspended in T0 medium. For each desired well of the AggreWell six-well plate, 2mL of cell suspension was prepared at a seeding density of 1.26 × 10^6^ cells per well. Cells in AggreWell plates were allowed to settle for 3 minutes before being spun at 200g for 5 min to promote aggregate formation. Cells were cultured in a hypoxia incubator at 5% CO2, 5% O2, at 37°C for the entire hemogenic endothelium induction culture phase.

2 ml of T1 medium was added to the 2mL of T0 media already in each well twenty-four hours after initial seeding in the AggreWell plates. The next day, after a total differentiation time of forty-two hours, medium was aspirated and replaced with 2mL per well of T1.75 medium. After seventy-two hours of total differentiation time, each well was topped up with an additional 2 ml of T1.75 medium. At a total differentiation time of ninety-six hours, medium was gently aspirated to avoid disturbing the aggregates and 2mL of T4 medium was then added to each well.

At a total differentiation time of 144 hours, 2 ml of T6 medium was added in addition to the 2mL of T4 media already in each well bringing the total volume to 4mL.

At a total differentiation time of 192 hours, media in each well was mixed thoroughly and aggregates were aspirated into 15mL or 50mL tubes and spun at 200g for 5 min (if the aggregate pellet is visibly loose, this time can be extended to 7 minutes). After aspirating the supernatant, the pelleted aggregates were resuspended in 3 ml of TrypLE supplemented with deoxyribonuclease I (MilliporeSigma, 260913-10MU) and incubated at 37°C for approximately 15 minutes. Cells were mixed approximately 30 times to achieve dissociation to a single cell suspension and TrypLE was diluted in HF buffer (Hanks’ balanced salt solution, Thermo Fisher Scientific, 14175103, + 2% FBS, Thermo Fisher Scientific, 12483020) and then cells were spun at 300xg for 5 minutes and washed in HF. Next the CD34-positive selection kit (Miltenyi Biotec, 130-046-702) was used to enrich for CD34+ cells which were cryopreserved using CryoStor CS10 (STEMCELL Technologies, 07930) following the manufacturer’s instructions.

#### Endothelial to hematopoietic transition culture

EHT media was prepared by combining complete StemPro34 (Thermo Fisher Scientific, 35050061) with 1x GlutaMAX (1x, Thermo Fisher Scientific, 35050061), 1-Thioglycerol (0.039 uL/mL, Sigma-Aldrich, M6145), Ascorbic Acid (50ug/mL, Sigma-Aldrich, A8960), Transferrin (150ug/mL, Millipore Sigma, 10652202001), bFGF (5ng/mL, PeproTech, 100-18B), VEGF (5ng/mL, R&D Systems, 293-VE), IL-6 (10ng/mL, R&D Systems, 206-IL), IL-11 (5ng/mL, R&D Systems, 218-IL), TPO (30ng/mL, R&D Systems, 288-TPN), IGF-I (25ng/mL, R&D Systems, 291-G1), SCF (50ng/mL, R&D Systems, 255-SC), IL-3 (10ng/mL, R & D Systems, 203-IL), ROCKi/Y-27632 (10uM, Sigma-Aldrich, Y0503), BMP4 (10 ng/mL, R&D Systems, 314-BP) and Flt3L (10ng/mL, R&D Systems, 308-FK). 96 well tissue culture-treated plates were coated overnight at 4°C with 50uL per well of a coating solution containing sterile phosphate-buffered saline (PBS), Fc-tagged VCAM1 (2.5 μg/ml; R&D Systems, 643-VM) and Fc-tagged DLL4 (15 μg/ml; Sino Biological, 10171-H02H). Prior to seeding, plates were pre-warmed to room temperature, coating solution was aspirated and wells were gently washed with 100uL/well sterile PBS.

Cryopreserved, post-CD34+ enriched cells were thawed and transferred dropwise into HF buffer pre-warmed to 37°C. Cells were pelleted by centrifugation at 300 x g for 5 minutes, supernatant was aspirated, and cells were re-suspended in freshly prepared, pre-warmed EHT media to a concentration of 1 × 105 cells/ml. 100uL of cell suspension (10,000 cells) were added to each well of the pre-coated and washed 96 well plate. EHT cultures were carried out for 7 days at 37°C, 5% CO2. Rounded hematopoietic cells were visible after 2 or 3 days and the non-adherent fraction of cells were collected on day 7 by gentle pipetting.

#### Progenitor T cell differentiation

For time course and lineage tracing experiments, the following “Cord-blood-optimized” serum-free media was used for T cell differentiation; Iscove’s modified Dulbecco’s medium (IMDM) with GlutaMAX basal medium (Thermo Fisher Scientific, 31980030) supplemented with 20% BIT 9500 serum substitute (STEMCELL Technologies, 09500), 60 μM ascorbic acid (Sigma-Aldrich, A8960), 24 μM 2-mercaptoethanol (Sigma-Aldrich, M3148), 0.05% low-density lipoprotein (STEMCELL Technologies, 02698), stem cell factor (SCF) (23.9 ng/ml; R&D Systems, 255-SC), Flt3L (8.7 ng/ml; R&D Systems, 308-FK), IL-3 (5.3 ng/ml; R&D Systems, 203-IL), CXCL12 (9.7 ng/ml; R&D Systems, 350-NS), IL-7 (10 ng/ml; R&D Systems, 207-IL), and TNFα (4.9 ng/ml; R&D Systems, 210-TA). In parallel work that took place during the course of this study, we developed an improved “PSC-optimized” progenitor T cell differentiation media which supports more efficient survival. We therefore used this media for CRISPR perturbation experiments, and their associated controls, to maximise viable cell recovery. The improved media comprises IMDM with GlutaMAX basal media (Thermo Fisher Scientific, 31980030) with 4% B27 minus Vitamin A (Gibco, 12587010), 60 µM Ascorbic acid (Sigma-Aldrich, A8960), 24µM 2-Mercaptoethanol, SCF (12.37 ng/ml), Flt3L (8.61 ng/ml), IL-3 (0.97 ng/ml; R&D Systems, 203-IL), CXCL12 (97.39 ng/ml), IL-7 (65.25 ng/ml), and TNFα (0.07 ng/ml).

Harvested cells from the EHT culture phase were pelleted, re-suspended in T cell differentiation media at a split ratio of 1:3 (unless otherwise specified), and transferred in 100uL volume per well on to 96 well plates freshly coated with DLL4 and VCAM1 as described above. Cells were incubated at 37°C, 5% CO2. The media was topped up with an additional 100uL of media per well (bringing the total volume of media to 200uL) after 3 or 4 days of progenitor T cell differentiation. Cultures proceeded for up to a maximum of 7 days.

### Single cell RNA sequencing time course

In order to mitigate confounding batch effects, we performed a reverse differentiation time course such that new differentiations were initiated every 24 hours for 14 consecutive days. Thus, samples representing each developmental time point could be collected at the same time for parallel capture

For each differentiation time point, we thawed an aliquot of the same cryopreserved stock of iPS11-derived hemogenic endothelial cells produced as described above. For each timepoint, at least 5 independent wells were seeded for downstream pooling. EHT cultures and progenitor T cell differentiations were performed as described above.

On the final day of the timecourse, all timepoint samples were harvested and an additional aliquot of the input population was thawed. For samples from the progenitor T cell differentiation stage, cells were harvested by pipetting, whereas for the EHT timepoints, non-adherent cells were harvested by pipetting before adherent cells were harvested using 30uL per well of 0.25%Trypsin EDTA (Thermo Fisher Scientific, 25200072). For the sample coresponding to the passage day, adherent and floating cells were sequenced separately.

Each sample was pelleted by centrifugation at 300x g for 5 minutes. Replicate wells were pooled together and washed with PBS. Samples were Fc blocked (BD, 564220) and labelled with zombie-UV viability dye (BioLegend, 423108) on ice in the dark for 20 minutes. Next, each sample was labelled with 0.1 ug of a cell hashing antibody (TotalSeq-B anti-human Hashtag antibodies 1 through 8, BioLegend). Cells were labelled for 30 minutes on ice in the dark. Next, samples were washed 3 times with HF buffer and counted prior to pooling. Samples were combined in to two pools, one for the EHT time points and another for the progenitor T cell differentiation time points. To enrich for Zombie-UV-negative viable cells, the EHT time point pool was subject to Fluorescent Activated Cell Sorting (FACS) using a MoFlo Astrios (Beckman Coulter). Cell pools were counted and loaded on the Chromium Controller (10x genomics) using the Chromium Single Cell 3’ Reagents kit (v3.1) and libraries were generated according to the manufacturer’s instructions. Libraries were sequenced on the NextSeq2000 (Read 1: 28bp Index1: 10bp Index 2: 10bp Read 2: 90bp)

### Single cell RNA sequencing time course analysis

#### scRNAseq pre-processing

FASTQ generation was done with the Cell Ranger function mkfastq. FASTQ trimming, alignment and counting with the hg19 reference was performed with the Cell Ranger count function.

Processing of the time course and lineage barcoded scRNAseq datasets was performed following the scanpy preprocessing tutorial (https://scanpy.readthedocs.io/en/stable/api.html). For the time course dataset, cells were captured in three reactions (pools A, C, and D) and were concatenated into a singular dataset before using the demultiplexed sample antibody counts from the CellRanger pipeline to filter empty droplets and doublets. Cells with fewer than 1000 genes, more than 20% mitochondrial reads, and more than 85000 reads were filtered to further account for doublets and dead/dying cells. Following filtering, the data was normalized to 10000 counts per cell, log-transformed, and highly variable genes were identified using scanpy. Principal components, neighbors, and a UMAP were then calculated. Cell cycle annotations were computed using scanpy and the provided gene set, and were subsequently regressed using the “regress_out” scanpy function. The processed time course dataset contained a final 7899 cells, 21888 genes.

The same preprocessing pipeline was followed with the lineage barcoded scRNAseq dataset. After processing the dataset contained 12642 cells and 18658 genes.

#### Anchor-based fetal-liver integration scoring on in vitro dataset

Anchor-based integration scores in Fig. 1f were calculated using a transfer-based approach implemented in the “Seurat” R package^37^. We used a publicly available in vivo reference dataset from fetal liver^32^. We used the “FindVariableFeatures” function (default settings) prior to PCA. Next next used the “FindTransferAnchors” function with 30 principal components and then applied the “TransferData” function to calculate primary cell type identity scores for each cell in the in vitro dataset.

#### Cell label transfer from fetal liver to in vitro dataset

To transfer cell labels from publicly available in vivo reference dataset of fetal liver^32^ to the generated in vitro dataset we trained an independent logistic regression model (LR, sklearn v0.24.0). The in vivo dataset was downsampled to 400 cells per cell type, to balance the cell number per cell type. Cell cycle genes were excluded from the modelling, to limit the influence of cell cycle. The in vivo dataset was split into 75/25 training and test set, and then identified 2,000 highly variable genes, normalized, log-transformed, scaled, subsequently. Hyper-parameter tuning was performed to maximize the accuracy of the model. Finally, the best prediction and probability per cell was calculated from the model.

#### Quality filters and alignment of in vivo and in vitro data

The generated in vitro data was integrated with publicly available in vivo fetal liver data ^32^. The filtered count matrices from Cell Ranger were integrated and analysed with scanpy (version 1.8.2), with the pipeline following their recommended standard practises. Briefly, for the in vivo dataset genes expressed in less than three cells and cells that have no annotation information, were excluded, while for in vitro dataset, genes expressed in less than three cells, cells expressing fewer than 500 genes and cells with more than 20% mitochondrial genes were excluded. Then, the normalized counts for each dataset was converted to log scaled and the object was transposed to gene space to identify cell cycling genes in a data-driven manner, as previously described^10,32,63^ from filtered count matrix. Principal component analysis (PCA), neighbour identification and Louvain clustering allowed us to identify the members of the gene cluster including known cycling genes. The members of the gene cluster were labeled as data-derived cell cycling genes, and discarded for downstream analysis.

Next, 5,000 highly variable genes from both datasets were identified using scanpy’s highly_variable_genes function with flavour set as default (seurat). The in vivo and in vitro data sets were integrated using the deep generative model single-cell variational inference (scVI, version 0.20.3). Number of layers in the scVI model was tuned manually to allow for better integration. The parameters used for modelling were 1 layer and 20 latent and the gene likelihood set to a negative binomial distribution with donor and experiment as covariates. The resulting latent representation of the data was used for calculating neighbourhood graph and uniform manifold approximation and projection (UMAP) representation.

#### Anchor-based thymus integration scoring on in vitro dataset

To validate cell type labelling, we used the transfer-learning approach from the R packages Seurat (version 4.3.0)^37^. Annotated RDS files were downloaded from Lavaert et al^9^. and Park et al^10^. through the Human Cell Atlas Data Portal. These and our data were loaded, and the top 2000 variable features identified using the variance stabilizing transformation option of ‘FindVariableFeatures’. Data was then scaled and centered using the ‘ScaleData’ function, and principal components identified with the ‘RunPCA’ function. Integration anchors were then identified between each reference dataset and ours based on the top 30 principal components using the ‘FindTransferAnchors’ function, and class label predictions made using the ‘TransferData’ function. For the Lavaert et al. dataset, we used the author_cell_type variable for the label transfer rather than cell_type variable as it had a higher degree of specificity. For both datasets, a global check of the prediction probabilities across our data for all cell types was first performed to ensure that the label transfer results were successful. For the Lavaert et al. dataset^9^, we examined how cells in our Pro-T Cell cluster compared over time to the T cell commitment status of the benchmarked data.

#### Cell label transfer from thymus to in vitro dataset by logistic regression

To transfer cell labels from publicly available in vivo reference dataset of Thymus (Park et al., 2020 and Suo et al., 2022) ^10,39^ to the generated in vitro dataset we trained an independent logistic regression model (LR, sklearn v0.24.0). The in vivo dataset was downsampled to maximum 1,500 cells per cell type, to balance the cell number per cell type. Cell cycle genes were excluded from the modelling, to limit the influence of cell cycle. The in vivo dataset was split into 75/25 training and test set, and then we identified 2,000 highly variable genes, normalized, log-transformed, and subsequently scaled. Hyper-parameter tuning was performed to maximize the accuracy of the model. Finally, the best prediction and probability per cell was calculated from the model, excluding the low quality cluster “high mito” from the in vivo datset.

#### PySCENIC analysis

The pySCENIC pipeline was used to compute regulons. Grnboost2 was employed for co-expression module inference and hg19 transcription factor motifs were used for direct transcription factor motif inference. The built-in area under curve metric quantified regulon activity in each cell.

#### Mutual Information vs Time

To determine the relationship between regulons and cell culture time, we used sklearn’s mutual information module. The mutual_info_classif function calculated mutual information for each regulon and time before ranking regulons based on their scores (https://scikit-learn.org/stable/modules/generated/sklearn.feature_selection.mutual_info_classif.html). This was done for each subpopulation of cells in the time course scRNAseq dataset to determine regulons with a strong relationship with the development of each lineage.

#### Hill model of regulon activity over time

Mutual information was computed as previously indicated between regulons and time for each cell type. Next, regulons that had a high mutual information with time were selected using the elbow method of feature selection. The resulting regulons were modelled over the time spent in culture using the scipy curve_fit function. Regulons were then sorted based on the variable k which represents the half maximal activity of the regulon to determine a point where regulons “switch” from a state of high activity to low activity or vice versa.

Hill function:

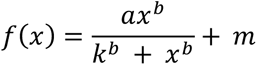

### Gene regulatory network modelling with IQCELL

#### Pseudo-time ordering

To establish a pseudo-time order for differentiating cells, we employ the Slingshot method ^64^. The Slingshot method generates a curve from the lower dimensional representation of the cells, and uses this to infer the relative sequence of the differentiating cells, beginning from the cells from day 7.

#### Normalizing cell density along the Pseudo-time axis

As the number of cells is unevenly distributed across the differentiation axis, we performed a down-sampling (to 1000 cells) of cells along the pseudo-time axis to obtain a more uniform distribution^65^. This approach can help mitigate any potential bias in GRN inference towards the densely populated areas in pseudo-time during subsequent steps.

#### Selecting active and dynamic regulons

First, we utilize the pySCENIC platform to deduce active regulons, which are groups of TFs along with their downstream targets^30^. Opting for regulons instead of solely TFs improves the selection process’s stability. We proceed by utilizing the active regulons to deduce a gene set for application in the GRN inference module. To select TFs, we consider their variability along the pseudo-time axis based on the gene dynamic score (GDS). We selected the TFs with the higher GDS. Top 20 Tfs were selected based on this step

#### Gene-gene interactions

Following the identification of the genes of interest, we proceed to compute both correlation (to calculate the type of interaction, +/-) and mutual information between pairs of genes. Firstly, mutual information is computed between each pair of genes. Subsequently, the top 30% of gene interactions are selected for downstream applications.

#### Correcting gene dropouts

TFs, or transcription factors, are particularly susceptible to dropout because they are typically expressed at lower levels and have fewer copies per cell compared to other genes. To address this issue, IQCELL utilizes MAGIC^45^ (with Knn value of 5) to compensate for the dropout effect and impute missing values in the scRNA-seq data. This improves the accuracy and reliability of downstream analyses.

#### Binarization

The process of binarizing genes involves converting gene expression values into binary ON/OFF states. This step is accomplished using the thresholding method based on K-means clustering (K=2).

#### Calculating gene hierarchy

Following the binarization of gene expression, we utilize the resulting data to compute the gene hierarchy. This hierarchy is determined by identifying the points at which genes transition from ON to OFF or OFF to ON. Genes that transition earlier in pseudo-time points are assigned higher positions in the gene hierarchy.

#### Obtaining functional Boolean GRN

Next, we employ the Z3 engine^66^ to identify the conceivable update regulations for each gene. Following that, we determine the optimal GRN according to the mutual information, selecting the GRN that maximizes the average mutual information of the GRN.

#### Simulating the GRN dynamics

To initiate the simulations, we begin by determining the starting state of the system, using the pseudo-time and binarized gene expression. Then, we simulate the Boolean GRN by using asynchronous Boolean(5) updates and generate resulting trajectories^67^.

#### Simulating Knockouts

To simulate systematic knockouts, we force the expression level of a gene to be turned off. After that, we use asynchronous Boolean simulation to calculate the resulting trajectory.

### Validating IQCELL predictions by polyclonal CRISPR perturbation

LentiCRISPRv2GFP (Addgene plasmid #82416) was using for sgRNA cloning as previously described. Briefly, the backbone was digested with Esp3I(NEB, R0734S), dephosphorylated and gel purified. Oligo inserts corresponding to each target sequence (See Table S1) were synthesized by IDT then annealed and phosphorylated prior to ligation to the backbone. Lentiviral particles were produced by co-transfecting the barcoding plasmid with the packaging and envelope plasmids pMD2G and pCMV-dR8.91 (provided by Andrew Ramos) into Lenti-X 293T cells (Takara, 632180) with Lipofectamine 2000 (Thermo Fisher, 11668027). The supernatant was concentrated using Lenti-X Concentrator (Takara, 631232) and cryopreserved for subsequent transduction. For knockout experiments in PSC-derived CD34+ cells, cells were seeded in EHT culture conditions as described above. 48 hours after seeding, wells were transduced with 4uL concentrated viral supernatant per well of a 96 well plate in 100 uL total volume of media. 24 hours later, transduced wells were topped up with an additional 100uL of fresh EHT media. One week after transduction, cells were passaged into progenitor T cell differentiation conditions at a split ratio of 1:4.

Flow cytometry analysis was performed at the end of the 7 day T cell progenitor differentiation stage using a CytoFLEX LX cytometer (Beckman), and data were compensated using single-stained controls in CytExpert v2.3. Gating and plotting were done with FlowJo v9.

### Umbilical cord blood culture and differentiation

Umbilical cord blood (UCB) samples were obtained from consenting donors in accordance with research ethics board approval. Samples were collected at BC Children’s Hospital. Mononuclear cells isolation was performed by density gradient centrifugation using). EasySep Human CD34+ Positive Selection Kit (Stemcell Technologies) was used to enrich for CD34+ cells to >90% purity.

For CRISPR perturbation experiments, cells thawed on to tissue culture-treated 96 well plates pre-coated as above, except using adjusted concentrations of 2µg/ml DLL4 and 1µg/ml VCAM-1. This is because UCB cells require lower levels of notch signalling to differentiate into progenitor T cells compared to PSC-derived cells. We seeded 2000 cells per well in 50uL of “cord-blood optimized” media (see Progenitor T cell differentiation section above for complete formulation. 24 hours after seeding, cells were transduced with 2uL per well of concentrated Lenti-CRISPR-v2-GFP supernatant (transduction efficiency kept <20% to promote single copy integration) and 1uL Vectofusin-1 (1X, Miltenyi Biotec, 130-111-163) diluted in 50 uL of cord-blood optimized media. 24 hours after transduction, an additional 100uL of media was added to each well. Four days post-transduction a half media change was performed. Six days post-transduction, cells were passaged at a 1 in 10 dilution on to plates freshly coated with DLL4 and VCAM1 as described above in fresh media. 3 days later, cells were collected for flow cytometry.

### Umbilical cord blood culture scRNA-seq analysis

FASTQ generation was done with the Cell Ranger function mkfastq. FASTQ trimming, alignment and counting with the hg19 reference was performed with the Cell Ranger count function.

The cord blood scRNA-seq dataset was processed with the scanpy package (https://scanpy.readthedocs.io/en/stable/). The dataset was filtered to remove cells with fewer than 2000 genes expressed, more than 12% mitochondrial reads, and with cells with greater than 40000 gene counts. The data was normalized to 10000 counts per cell and was log1p transformed before determining highly variable genes. Principal component analysis was performed, and a neighbourhood graph was computed before computing a UMAP embedding. Leiden clustering was performed and differentially expressed genes computed. The final scRNA-seq dataset consisted of 6322 cells and 20598 genes.

### Lineage barcoding library cloning

An oligonucleotide containing 16 randomized bases was synthesized by IDT (Lin_Barcode_insert_v1: AGCTAGCTTGGAGACTGCGTATGATNNNNCTNNNNACNNNNTCNNNNGTGCTGATACCGTTATTAACATATGACAACTCAATTAAACACTGACCGGTGGA). This oligo was PCR amplified using the primers Lin_Barcode_cloning_F1 (GTCTCTCAGCTAGCTTGGAGACTGCGTATG) and Lin_Barcode_cloning_R1(GATGTCCACCGGTCAGTGTTTAATTGAG). 17 PCR cycles were performed using Phusion High-Fidelity PCR Master Mix with GC Buffer (NEB, M0532) with primers and template oligo each at a concentration of 10uM. Following an initial denaturing step of 98°C for 30 seconds, each cycle comprised melting (98°C for 12 seconds), annealing (63°C for 15 seconds) and extension (72°C for 15 seconds). A final 5-minute extension step was performed at 72°C. This PCR was performed in triplicate to mitigate amplification bias and the product was purified using the MiniElute PCR purification kit (Qiagen, 28004). The insert was then digested with AgeI-HF (NEB, R3552) and NheI-HF (NEB, R3131) and re-purified as above.

The destination vector (EGFP-miSFIT-SCR, derived from Addgene plasmid #124690) was digested with AgeI-HF (NEB, R3552) and NheI-HF (NEB, R3131), dephosphorylated with Antarctic Phosphatase (NEB M0289) and column purified (Qiagen, 28104). The digested and purified inset and vector were ligated at a ratio of 5:1 using T4 DNA ligase (NEB, M0202) overnight at 18°C. The resulting ligation product was column purified, eluted in nuclease free water and transformed into NEB 10-beta electrocompetent E. coli (NEB, C3020K). 4 electroporations of 400ng ligation product each into 50uL of bacteria were performed according to the manufacturer’s instructions. We plated a 1/30,000 dilution of the pooled electroporation to estimate a total recovery of 1.02 x 10^6^ colonies. To verify the library complexity, a PCR was performed with the primers Complexity_seq_F1: agcttggagacTGCGTATGAT and Complexity_Seq_R1: cagccataccacatttgtagagg. Following adapter ligation and indexing, the amplicon was sequenced on the Illumina Next-Seq. We named the resulting vector library EGFP_miSFIT_Lineage_BC_lib (see supplementary workbook 1 for sequence).

Lentivirus was generated by co-transfecting the barcoding plasmid with the packaging and envelope plasmids pMD2G and pCMV-dR8.91 (provided by Andrew Ramos) into Lenti-X 293T cells (Takara, 632180) using Lipofectamine 2000 (Thermo Fisher, 11668027). Viral supernatant was concentrated using Lenti-X Concentrator (Takara, 631232).

### Simultaneous recovery of transcriptomes and clonal identity

#### Experimental design

In vitro differentiations from cryopreserved CD34+ cells were initiated as described above, with the modification that 7500 cells were seeded on to one well of 96 well plate for transduction and sampling while additional “proxy” wells were seeded at the same density for flow cytometry analysis and estimating the cell count in the experimental well. Three days after seeding, cells were transduced with 3.75uL per well of the concentrated barcoding virus. Twenty-four hours later, wells the cells were fed with an additional 100uL per well of EHT media. Five days after seeding (48 hours after transduction), the well was mixed by gentle pipetting and half of the total volume was sampled for scRNA sequencing (Chromium Single Cell 3’ Reagents kit v3.1). One week following the initiation of the differentiation culture (4 days after transduction), the remaining cells were collected and passaged 1:2 on to plates coated with DLL4 and VCAM1 in EHT media as described above. Three days later, remaining cells were harvested, counted and a sub-sample of 16,500 cells were taken for scRNA sequencing.

#### Sequencing library generation

Gene expression libraries were generated using the Chromium Single Cell 3’ Reagents kit (v3.1) following the manufacturer’s instructions. To simultaneously capture and sequence lineage barcodes and 10x cell barcodes, a targeted amplicon library was generated by PCR from the standard gene expression library cDNA using Extract-N-Amp PCR ReadyMix 2x (Sigma, E3004). 50uL PCR reactions were prepared by combining 25uL ReadyMix with 2.5 uL of a 10uM stock of each primer (10x Partial Read 1: CTACACGACGCTCTTCCGATCT and a custom primer Lineage_BC_seq_Rev: TGACTGGAGTTCAGACGTGTGCTCTTCCGATCTCTAGCTTGGAGACTGCGTATG), 5ng cDNA template and nuclease-free water up to 50uL total. PCRs were performed for 23 cycles. Following an initial denaturing step of 94°C for 8 minutes, each cycle comprised melting (94°C for 30 seconds), annealing (50.7°C for 30 seconds) and extension (72°C for 30 seconds). A final 5-minute extension step was performed at 72°C. Next, the products were purified and size selected by SPRI clean-up (SPRISelect, B23319 Beckman Coulter) using a double size selection (0.75x for left side/ 0.65x for right side).

The purified libraries were indexed by PCR using Extract-N-Amp PCR ReadyMix 2x (Sigma, E3004) and sample-specific i5 primers (10x genomics PN2000095) and i7 primers (10x genomics Single Index Kit T Set A PN2000240). After initial denaturation at 98°C for 45 seconds, 14 PCR cycles were performed with a 20 second melt at 98°C, 30 second annealing at 54°C, and a 20 second extension at 72°C. A final extension step at 72°C for 1 minute was performed. The final amplicon was purified using SPRI beads (1.0x).

#### Extracting lineage barcode reads from FASTQ files

To extract and process lineage barcode reads from the targeted barcode sequencing we first combined read1 and read2 FASTQ files using the UMI-tools (v1.1.1) extract function (https://umi-tools.readthedocs.io/en/latest/). From the subsequent combined FASTQ files the grep function was used to extract reads that contained the pattern of our lineage barcode construct (ATNNNNCTNNNNACNNNNTCNNNNGT). To confirm that each extracted read matched sufficiently to the lineage barcode construct sequence, the Biopython’s pairwise2 local alignment tool was used to calculate alignment scores and filter out poorly aligned reads (https://biopython.org/docs/1.75/api/Bio.pairwise2.html).

Given that many of the reads contained single nucleotide sequencing errors, we wanted to ensure that reads with the same UMI but different lineage barcodes were not counted as separate lineage barcode reads. Reads with the same UMI but with lineage barcode sequences within one unit of Levenshtein distance were combined into a single UMI-lineage barcode combination by selecting the UMI-lineage combination with the most reads. Reads with the same UMI but lineage barcode sequences >1 unit Levenshtein distance apart were discarded (https://pypi.org/project/python-Levenshtein/).

To further account for single nucleotide sequencing errors, we created a set of “consensus barcodes” of lineage barcodes that had more than one count in the dataset. If another lineage barcode was only found once but was a single nucleotide error from matching a consensus barcode the barcode was matched to the consensus barcode. If a lineage barcode only appeared once, it was discarded. Barcode reads were only kept if they were associated with a cell barcode in the final scRNA-seq data set. Additionally, if a lineage barcode only appeared with one cell barcode in the final scRNA-seq dataset, the lineage barcode was discarded. We ensured that there were a minimum of two barcode counts (each with a different UMI) for a cell barcode-lineage barcode combination. The final filtering step determined the lineage barcode with the highest number of counts for each cell barcode and removed barcodes that had fewer than 50% of the counts of the most abundant lineage barcode for that cell.

#### Clone construction and processing

After extracting the lineage barcode-cell barcode counts we represented our clones using graphs using the python package networkx (https://networkx.org/). Graphs were constructed such that a node in the graph represented a cell. This cell was defined as a collection of reads that shared a singular cell barcode sequence. Edges in the graph were used to designate a lineage barcode sequence that was shared between two cells (i.e. reads of CBa-LB1 and CBb-LB1 creates an edge between CBa and CBb). After inputting all the CB-LBC reads into graph format the data was separated into independent subgraphs that represented the unprocessed “clones”. To correct for noise in the data and create high confidence clones we devised a processing algorithm for our clones. The processing algorithm utilised the degree of nodes in each subgraph and checked for possible subclones connected by artefact using a flow-based minimum cut algorithm.

Clones were processed through the following steps:

a. Flow-based minimum edge cut algorithm removes edges from the graph. The result of the edge removal was considered a successful separation of two subclones if the result met the following criteria.

i. The original clone was separated into subclones by the cut
ii. Each subclone has at least two nodes (cells)
iii. Each subclone contains at least 25% of the original graphs nodes (as to avoid creating false clones that should simply be pruned and discarded)

If the criteria were met, then the new subclones would go through further processing starting with step c). If not, move to step b) with the original clone.

b. Flow-based minimum node cut algorithm removes a node from the graph. The result of the edge removal was considered a successful separation of two subclones if the result met the following criteria.

iv. The original clone was separated into subclones by the cut
v. Each subclone has at least two nodes (cells)
vi. Each subclone contains at least 25% of the original graph’s nodes (as to avoid creating false clones that should simply be pruned and discarded)

If the criteria were met, then the new subclones would go through further processing.

c. Nodes (cells) were filtered from the clone based on the node degree. Nodes were removed if the node degree was <=55% that of the max degree node. NOTE: If a break was created by this filtering process, subsequent filtering was performed on the new subclones.
d. Cutting attempt: repeat step a and b.
e. Filter by clone connectivity (only used because of the possibility of multiple edges present between two cells).
f. Cutting attempt: repeat step a and b.

#### Clone Type Assignment

Clones were assigned to a certain “clone type” based on their day 10 cell population composition. For a given clone, if all of the cells from the day 10 time point were the same cell type, then the clone type would be assigned as that cell type. If a clone had more than one cell type at day 10, then the clone would be labelled as “multitype”.

#### Integrating the lineage barcoding dataset with the time-course

The lineage barcoded dataset (of cells at days 5 and 10) was integrated into the day 5 and 10 samples of the time-course dataset using regression with Seurat v3. Integration was performed for the samples at day 5 and 10 separately as this was found to produce better results than computing the regression with both together. Choosing anchors in the time-course samples only allowed the integrated day 5 and day 10 barcoded samples to be considered as samples in the full time-course without regression of the samples from other timepoints being required. The set of genes in the integrated dataset was reduced to the 18545 genes common to the two constituent datasets. Combat, mnnCorrect and scanorama were also tested as methods for dataset integration, with Seurat v3 found to produce the best results. The Seurat v3 standard workflow was run with default parameters with the exception of using log normalization for identifying the anchors and performing the integration, and setting the number of neighbours for weighting anchors to 50. Methods were assessed by whether cells in each cell type annotated in the raw datasets retained its cluster structure and whether the regression adjusted each cell type near to the position of corresponding type(s) in the anchored dataset. Assessment was performed by visual inspection of UMAP and PCA (2 components) of the integrated dataset.

### Clone dataset analysis

#### Determining principal component distance between clones

To confirm that cells within a clone demonstrated transcriptomic similarity, we compared principal component (PC) distance between cells within clones in our scRNAseq dataset and PC distance between cells in randomly generated clones. First, null clones were created by scrambling our clones. Subsequently, for both the real and simulated clones we found the mean euclidean PC distance (based on the 50 computed principal components) between the cells in each one of the clones. The distribution of mean euclidean PC distances for the real clones was compared to that of the null clones.

#### Oversampling Two Cell Combinations within Clones

In order to investigate the developmental relationships between the lineages in the scRNAseq dataset we found the relative probability that cell types co-occurred in a clone. First, 1500 simulated clone datasets were generated by assigning cells from our Day 10 clones to random clones. The simulated clone datasets followed the same cell per clone distribution as the real dataset. Then for the real clones and each of the 1500 simulated clone datasets, 20000 cell type comparisons were performed by selecting a cell at random, then a second cell belonging to the same clone at random before comparing the cell types of the two cells. The frequency of cell type combinations in each dataset was determined. The results from the real clones were normalized to those from the simulated clones. The simulated clones acted as a null distribution of clones that would occur if there was no bias in the likelihood that two cell types arise from the same ancestor. From this, we were able to determine the developmental relationship between lineages in the dataset.

### Mathematical trajectory inference

LineageOT is an integrated framework that reconstructs lineage and computes trajectory inference ^59^, which is an extension of WaddingtonOT^61^. The model of Optimal Transport is used to reconstruct trajectories based on time-course data of cell states and lineage information. The lineage data allows for greater accuracy and fewer time points when we explore the complex transition among cell states.

Since lineage information was available at time points 5 and 10, we used LineageOT to compute couplings between (4, 5) and (9, 10). We also used WaddingtonOT to compute couplings between other successive time-points and composed couplings.

The method requires as input an estimate of growth. Hence, in the following section, we will describe how to perform the estimation.

#### Growth rate estimation

We apply the logistic functions based on gene signature scores to estimate growth rates before day 8. After day 8, when the MPP/HSC population has dissipated, we assume that subsequent changes in cell abundance are due to growth and death rather than differentiation and use changes in proportions of cell types to estimate growth rates including and after day 8. After day 8, we use the assumption that descendants only come from the cells in the same cell type from the previous time point, and changes in cell type abundance is a more direct predictor of relative aggregated growth and death than gene expression signatures.

#### Gene signature score

The cell types of the annotated data include Progenitor cells, Myeloid cells, Erythroid cells, Mast cells, Endothelial cells, and ProT cells. The G2/M marker genes and the apoptosis genes were loaded from external gene set files^68^. The method gene_set_scores from WaddingtonOT then calculates the scores for each cell on apoptosis and proliferation with the gene sets described above as the input (https://broadinstitute.github.io/wot/cli_documentation/).

#### Logistic function

For an initial estimate of growth, we used gene signature scores, computed as follows. Logistic functions are used to estimate the growth rates for the cells before day 8 in both multi-time lineageOT (MT-LOT) and WaddingtonOT methods. The birth rate is estimated by 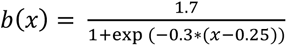, where *x* is the G2M observation. The death rate is calculated by 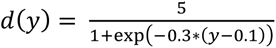, where *y* is the cell apoptosis signature score. The parameter values of these logistic functions were tuned to set the maximal and minimal values of growth rates. Eventually, the growth rate is approximated by *g*(*x*,*y*) = exp(*b*(*x*) – *d*(*y*) if the time difference is one day. The parameter values of these logistic functions were tuned to set the maximal and minimal values of growth rates.

#### Estimation with proportions

For the time points after and including day 8, the cell growth rates are estimated by the changes in the number of observations for each cell type. On days 9, 10, and 11, the proportions of the number of observations for each cell type within the total number of observations are calculated. The averages of two ratios, the proportion between day 10 and day 9 as well as the proportion between day 11 and day 10, are obtained to be the estimated growth rates. Relative growth rates were used as the marginals to compute the lineage coupling between the given time points.

#### MT-LOT transition map

We use the data obtained from the lineage barcoding experiment to generate transition maps using lineageOT with multi-time information for the time pairs (4, 5) and (9, 10)^62^. The following filter was applied for each time pair as the preprocessing step.

First, given the time pair (t1, t2), we filter for the data on day t1, day t2, or day 5. Then, the clones at day t2 and day 5, which have no multi-cell (at least two cells at day t2) or no multi-time (at least one cell at both day 5 and the second time point) lineage barcode information, would be filtered out. The cells on day t1, the cells on day t2 with barcode, and the cells on day 5 with barcode formed a subset of the original annotated data. This new set was then fed into lineageOT to fit a lineage tree from static lineage tracing (https://lineageot.readthedocs.io/en/latest/).

Next, we estimated the growth rates either by the logistic functions or the change in proportions of the observations for each type. The detailed method will be explained in the Growth rate estimation section. The marginal distributions on day t1 and day t2 could be calculated with the estimations, and they eventually served as the inputs to fit a completed lineageOT coupling. To compare the performance of MT-LOT with other methods (such as MT-Cospar or WOT), we concatenate two transition maps generated by MT-LOT with the maps generated by WOT. Similar filters were applied for data on day 5 and day 10 to ensure consistency in the number of observations between the two methods.

#### WOT transition map

MT-LOT method was only applied for time pairs (4, 5) and (9, 10). For the remaining time pairs, the package WaddingtonOT computed the transport maps without applying specific multicellular filters.

#### CoSpar analysis

CoSpar (Wang et al., 2022) was used to provide a prediction of the trajectory coupling between the day 5 and day 10 cells. Recall that day 5 and 10 are the two sample times at which lineage barcodes were detected. The default sub-method of CoSpar was used as it can make use of the data (lineage barcodes observed over two time points and gene expression) available from the barcoding experiment. As a lineage barcode was not detected for a subset of the cells sampled at these time points, CoSpar was run with the parameter extend_Tmap_space=True to obtain a trajectory coupling which included all cells sampled. Otherwise the default parameters of the algorithm were used to obtain the results, after tuning experiments (random sampling) determined no improvement from changing the values (defaults were: smooth_array=[15, 10, 5], CoSpar_KNN=20, sparsity_threshold=0.1, intraclone_threshold=0.05, normalization_mode=1, trunca_threshold=[0.001, 0.01]). The other two sub-methods of CoSpar (one using gene expression only, one using lineage barcodes observed at a single time and gene expression) were also tested but were not considered in our final analysis as the results were judged to be less accurate. This was determined by evaluating agreement with the developmental pathways present in the raw lineage barcode data and the biologically plausibility of the distributions of the fate probability predictions derived from the trajectory couplings.

## Ethics declaration

Y.S.M., P.W.Z., and J.M.E. are named inventors on patents for T cell differentiation technology. P.W.Z. is a cofounder of Notch Therapeutics. P.W.Z. and Y.S.M. are co-founders of Apiary Therapeutics. P.W.Z., Y.S.M., and J.M.E. consult for cell therapy companies. The authors declare no other competing interests.

## Acknowledgments

Work in P.W.Z’s lab was supported by Notch Therapeutics, the Stem Cell Network, the Canadian Institute for Health Research and the Wellcome Leap Human Organs, Physiology, and Engineering (HOPE) program. Y.S.M. was supported by a Michael Smith Foundation for Health Research Trainee award and the Canadian Institute for Health Research (CIHR) Banting Fellowship and is supported by the CancerCare Manitoba Foundation. M.C.M. was funded by a Natural Sciences and Engineering Research Council of Canada (NSERC) USRA and CBR summer award. J.M.E. was funded by a NSERC Canadian Graduate Scholarship and Zymeworks–Michael Smith Laboratories Fellowship in Advanced Protein Engineering. E.L.C. is a CIHR Vanier scholar. N.W. acknowledges the Allen Distinguished Investigators Grant. R.V-B. is supported by EMBO Postdoctoral Fellowship (ALTF 737-2021). R.V-T is supported by Wellcome Sanger core funding (220540/Z/20/A). A.F. was supported by a research fellowship from Royal Commission for the Exhibition of 1851.

