## Supplementary Figures for "Time- and lineage-resolved transcriptional profiling uncovers gene expression programs and clonal relationships that underlie human T lineage specification"

### Supplementary information

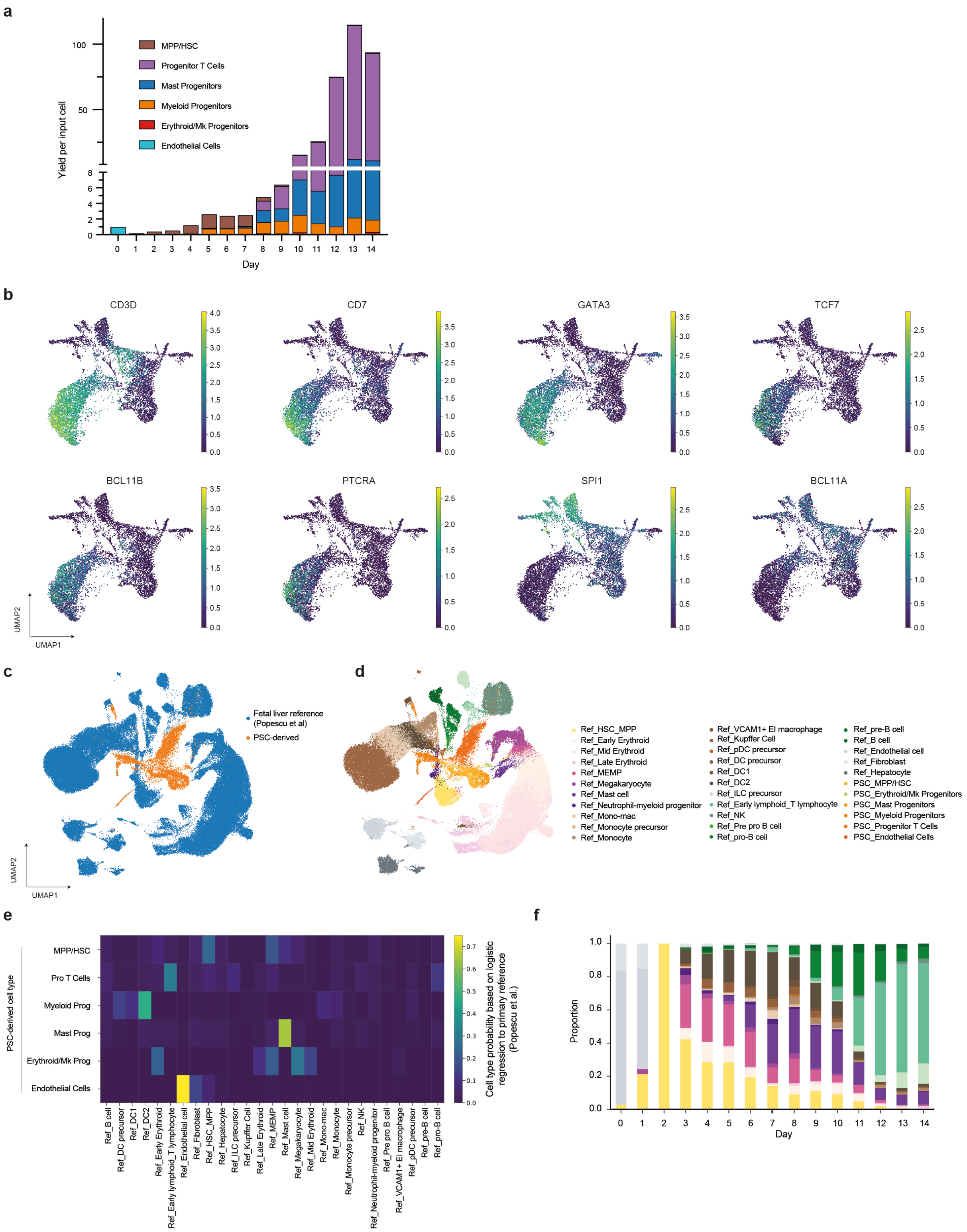

**Supplementary Figure S1: Pluripotent stem cell-derived endothelial cells undergo endothelial to hematopoietic transition and produce diverse blood cell types. a.)** PSC-derived cells enriched for CD34 expression (day 0) were seeded into culture conditions that support EHT for 7 days and transferred to T cell progenitor differentiation conditions for an additional 7 days. Cells at each time point were counted prior to single cell capture and sequencing. The abundance of each cell type at each time point is displayed. **b.)** Expression of a subset of T-lineage associated genes overlaid on Uniform manifold approximation and projection (UMAP) projections. **c.)** UMAP of scRNA-seq of in vitro timeseries data integrated with in vivo hematopoietic cells from the fetal liver (Popescu et al) coloured by dataset. **d.)** UMAP of scRNAseq in vitro timeseries integrated with in vivo hematopoietic cells from the fetal liver coloured by cell states. **e.)** Heatmap showing the predicted probabilities that PSC-derived in vitro differentiated cells map to primary cell types according to our logistic regression model. (0: low, 1: high probabilities). We built and trained the logistic regression classifier on Popescu et al., 2019 dataset and projected it onto the in vitro timeseries. In vitro cell types are represented in rows and in vivo cell types in columns. **f.)** PSC-derived cells were annotated based on their highest-probability primary cell type equivalent. Bar plot shows cell type proportions per timepoint.

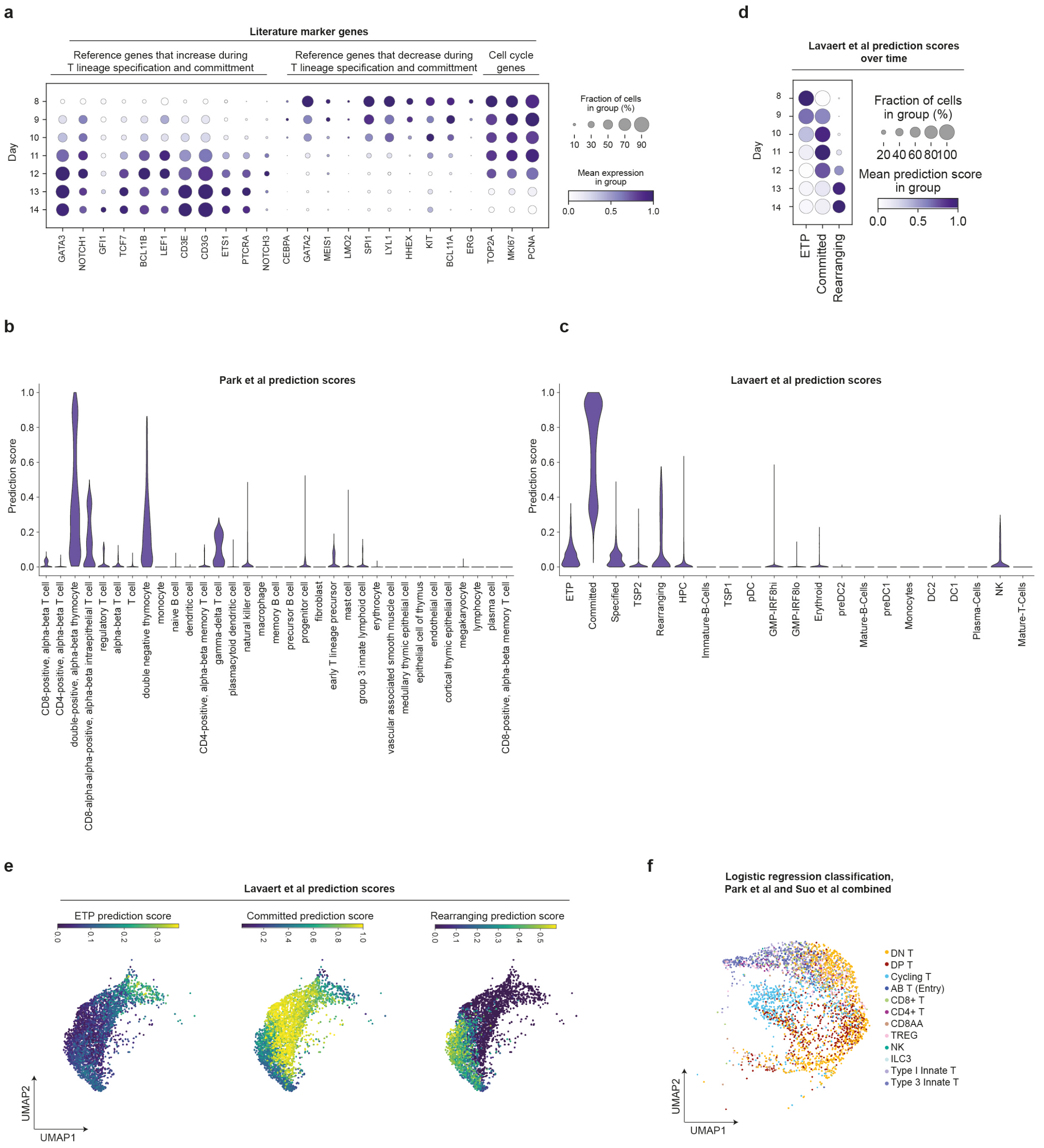


**Supplementary Figure S2: Pluripotent stem cell-derived T cell progenitors express a transcriptional program corresponding to primary committed T cell progenitors. a.)** Expression of marker genes known to be associated with T cell specification and commitment. Expression of representative cell cycle genes indicate that cells exit the cell cycle as they progress down the T lineage, consistent with primary T cell development. **b.)** Transcriptome-wide anchor-based integration was applied to PSC-derived progenitor T cells to calculate prediction probability scores for primary cell types using a human thymus reference dataset (Park et al). **c.)** Anchor-based integration as in (b) using a second primary reference dataset from Lavaert et al. This dataset contains a higher proportion of double-negative T cell progenitors and cell type labels with increased resolution. **d.)** Dotplot displaying cell type prediction scores over time for PSC-derived progenitor T cells. **e.)** Uniform Manifold Approximation and Projection (UMAP) of primary cell type prediction scores projected on to PSC-derived T cell progenitors. **f.)** A second independent integration method verifies the T-lineage identity of our PSC-derived T cell progenitors. UMAP (Uniform Manifold Approximation and Projection) of PSC-derived progenitor T cells labelled using a logistic regression classifier with the annotations from Suo et al.

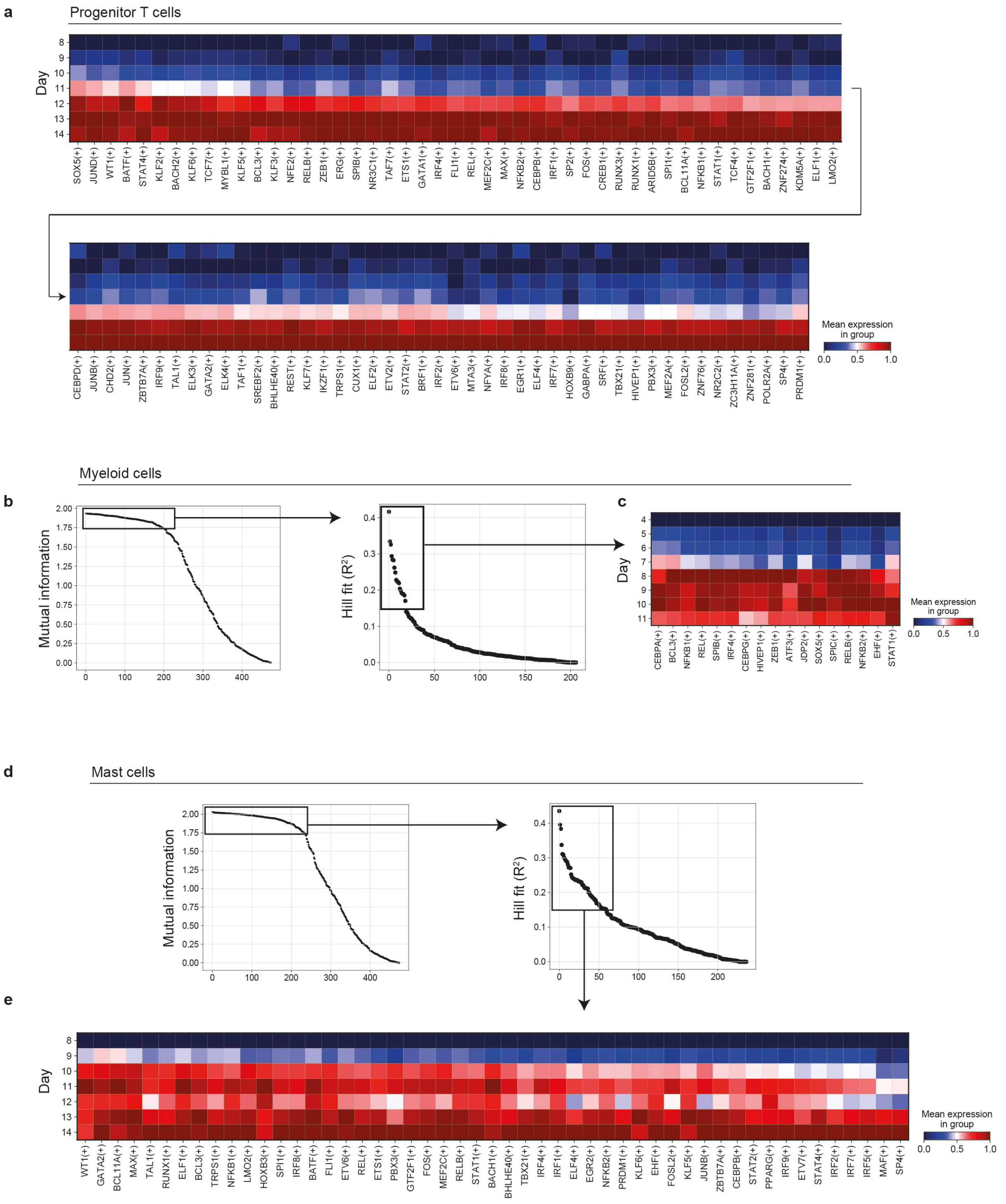


**Supplementary Figure S3: Temporal dynamics of transcription factor activity during T cell and mast cell differentiation. a.)** Transcription factor programs that turn on during T cell differentiation, as determined using the unbiased analysis pipeline in Fig 2a, are plotted over time. **b.)** Using the same analysis pipeline outlined in Fig 2a, we identified transcription factor programs that turn on during myeloid differentiation. **c.)** Activity of the transcription factors identified in (b) are plotted over time for myeloid progenitor cells. **d.)** Using the same analysis pipeline outlined in Fig 2a, we identified transcription factor programs that turn on during mast cell differentiation. **e.)** Activity of the transcription factors identified in (d) are plotted over time for mast cell progenitors.

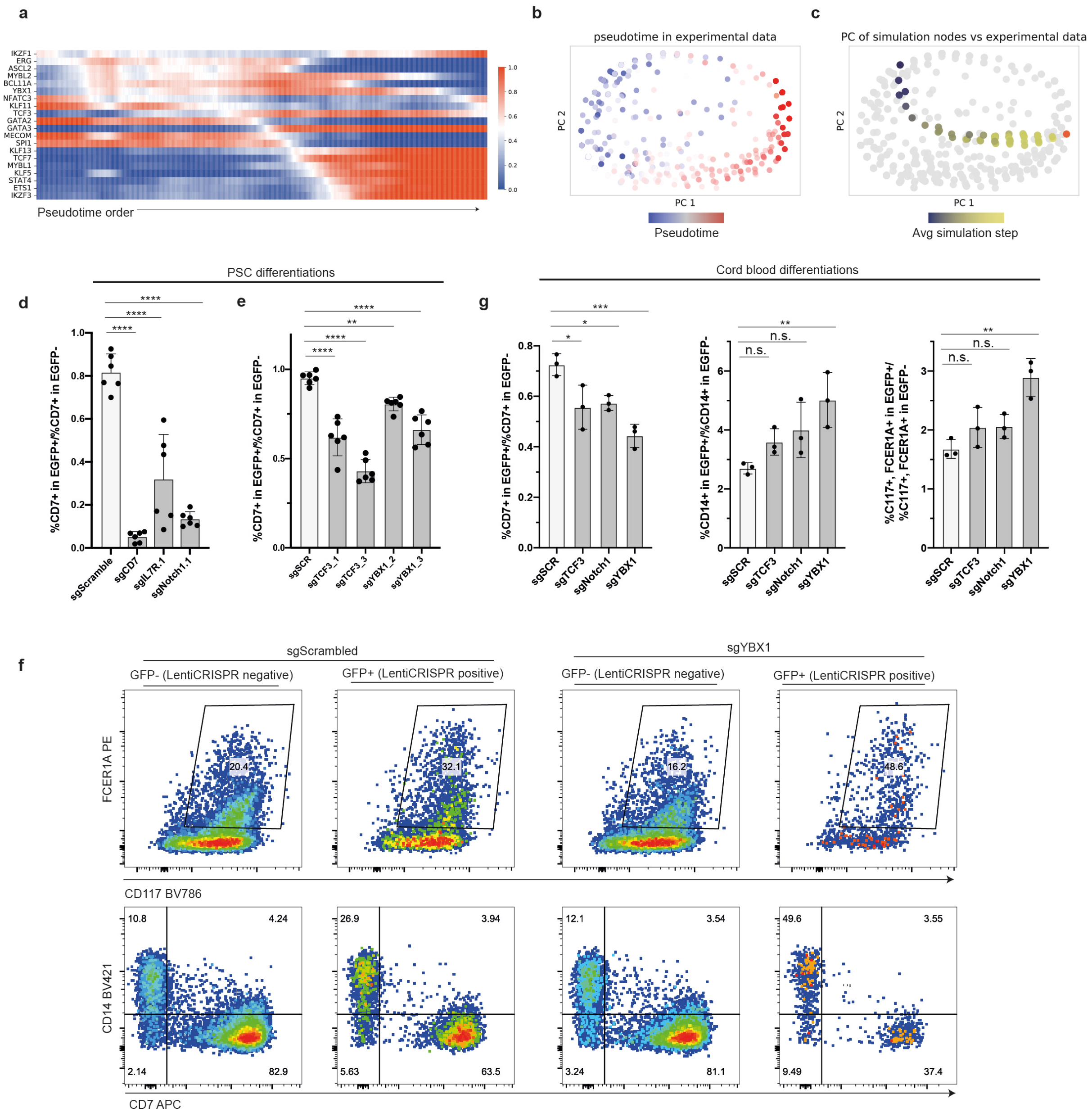


**Supplementary Figure S4: Predicting and validating a role for YBX1 in T cell specification. a.)** Smoothed gene expression matrix for cells along the T cell differentiation trajectory. Genes shown are components of the gene regulatory network predicted by IQCELL. Average of binarized expression values were calculated over a pseudo-time window. High expression is indicated in red and low expression is shown in blue. **b.)** Principle component (PC) plot for binarized experimental single cell expression data along the T cell differentiation trajectory. Cells are coloured by pseudotime corresponding to the order in (a) with early times shown in blue and later time points in red. **c.)** PC plot for IQCELL simulated developmental trajectory overlaid on top of the real data points in (b). An identified steady-state attractor is shown in red. Simulated points are coloured by the average step in the simulation with blue points corresponding to early steps in simulated development and gold points corresponding to later steps in the simulation. **d.)** Quantification of CD7+ progenitor T cell output from PSC-derived cells transduced with Cas9 and the indicated sgRNA for positive control sgRNAs targeting genes known to be important for early T cell development. **e.)** Quantification of CD7+ progenitor T cell output from PSC-derived cells transduced with Cas9 and the indicated sgRNAs targeting YBX1 and TCF3. **f.)** Representative flow cytometry analysis of CD34+ cord blood cells transduced with Cas9,

EGFP and the indicated sgRNA and then differentiated. gating strategy used to identify FCER1A+, CD117+ mast cell progenitors, CD7+ T cell progenitors and CD14+ myeloid progenitors. **g.)** Quantification of differentiation outcomes for cord blood transduced with Cas9, EGFP and the indicated sgRNA. In (d-g), error bars indicate standard deviation. **** = adjusted $P < 0.0001$, ** = adjusted $P < 0.01$, * = adjusted $P < 0.05$, Dunnett's multiple comparisons test.

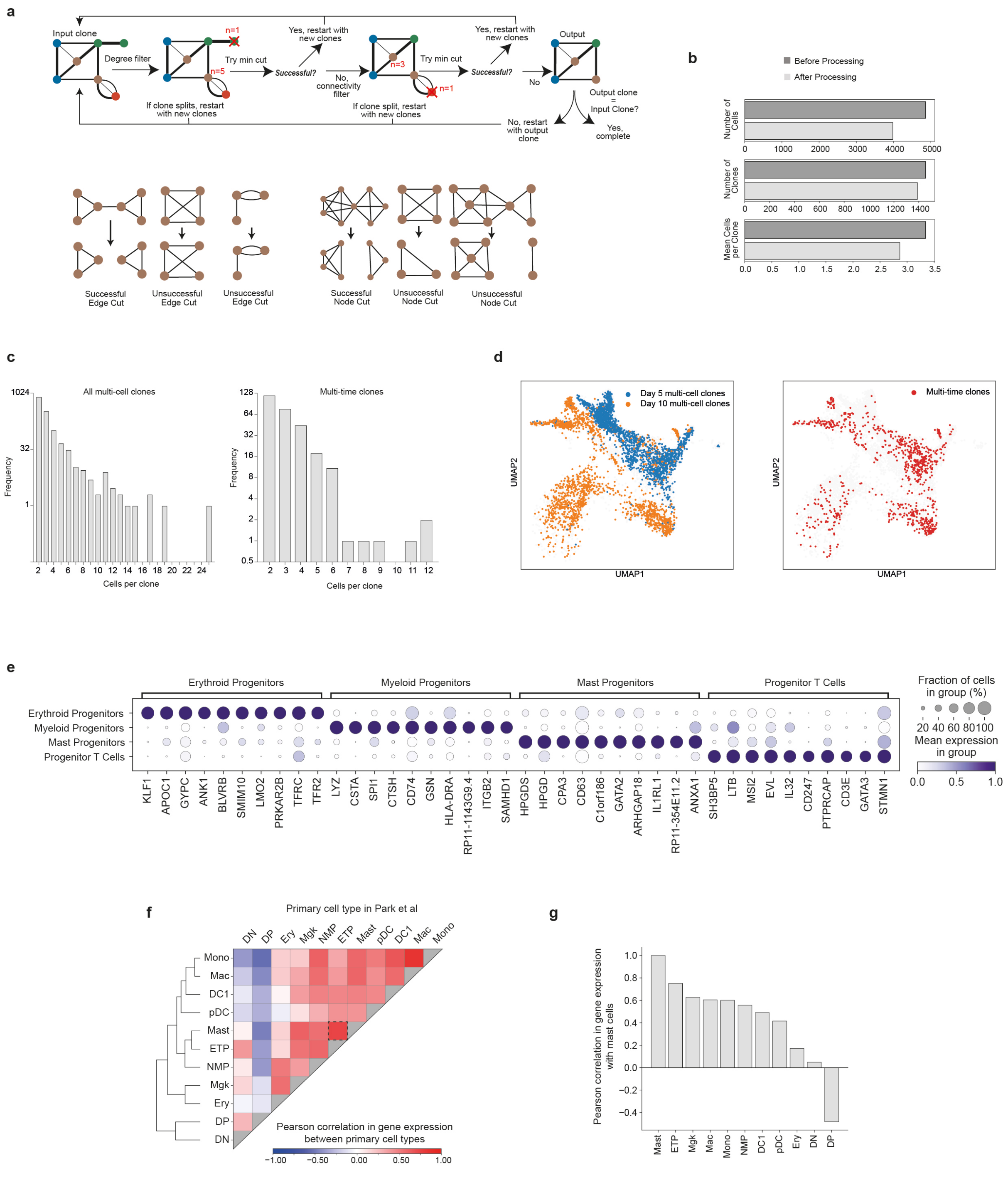


**Supplementary Figure S5: Transcribed lineage barcoding tracks the clonal relationships between hematopoietic cells differentiated from PSCs and cord blood in vitro. a.)** Top: Graph-based filtering workflow to identify high-confidence clones. Bottom: examples of successful and unsuccessful edge and node cuts. Nodes represent cells and edges represent shared lineage barcodes. **b.)** Metrics for clone assignment for PSC-derived cells before and after graph-based filtering. **c.)** Distribution of clone sizes for PSC-derived multi-cell and multi-time clones following graph-based clone filtration. **d.)** Uniform manifold approximation and projection (UMAP) projections of multi-cell and multi-time PSC-derived clones. **e.)** Top 10 differentially expressed genes associated with each cell type identified at day 7 of cord blood differentiation. **f.)** Pearson correlation analysis comparing transcriptional similarity between cell types found in the primary human thymus from Park et al.

Dendrogram shows results of hierarchical clustering based on gene expression **g.)** Bar graph of Pearson correlation coefficients between mast cells and each other hematopoietic cell type in the Park et al thymus reference atlas.

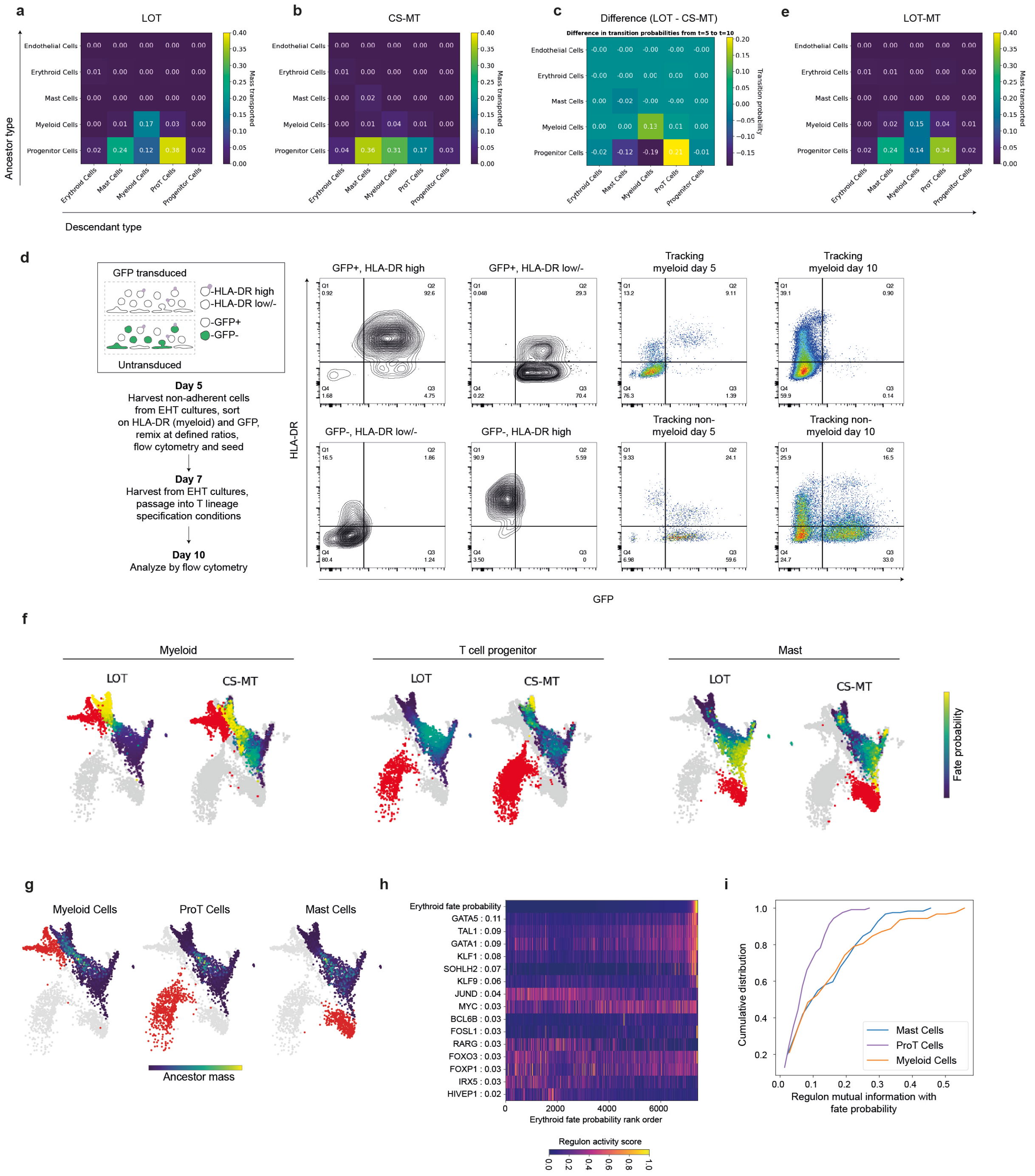


**Supplementary Figure S6: Mathematical trajectory inference of cell fate transitions during T cell differentiation from PSCs. a,b.)** Transition tables predicted by Lineage Optimal Transport (LOT) (a) and CoSpar-Multi-time (CS-MT) (b). Values indicated the predicted cellular mass transported between day 5 ancestor populations and day 10 descendant populations. **c.)** Differences between trajectory inference methods in the predicted mass transfer was calculated by subtracting the CS-MT values from LOT. **d.)** Experimental design and flow cytometry results for a fate tracking experiment to determine whether day 10 myeloid cell primarily arise from pre-existing day 5 myeloid cells or from day 5 non-myeloid cells. Results indicate that LOT underestimates the contribution of day 5 progenitor cells to day 10 myeloid cells, which motivated us to implement multi-time LOT (MT-LOT). **e.)** Transition table predicted by MT-LOT. **f.)** UMAP projections of day 5 cells from the lineage barcoding experiment coloured by the probability that they are fated to the specified downstream lineage at day 10 calculated using LOT and CS-MT. Day 10 cells for the indicated lineage are coloured red. **g.)** UMAP projections of day 5 cells from the lineage barcoding experiment coloured by their predicted contribution (ancestor mass) to the specified downstream lineage at day 10 calculated using MT-LOT. Day 10 cells for the indicated lineage are coloured red. **h.)** Transcription factor programs associated with erythroid fate. TFs were identified by calculating TF activity scores using SCENIC and calculating mutual information between SCENIC TF activity and MT-LOT fate probabilities. Numbers following the TF name indicate the mutual information score. Each column corresponds to a cell and cells were ordered from left-to-right based on increasing probability that the cell is fated for the specified lineage. **i.)** Cumulative distribution plot of mutual information scores between transcription factor activity (SCENIC regulon scores) and cell fate probabilities (scored by MT-LOT) for cells that have not yet been annotated as being part of the mast, myeloid or progenitor T cell cluster.
